# Long-read whole-genome sequencing offers novel insights into the biology, stress adaptation, and virulence of neurotropic dematiaceous fungi associated with primary cerebral phaeohyphomycosis

**DOI:** 10.1101/2025.01.02.631048

**Authors:** Arghadip Samaddar, Jenevi Margaret Mendonsa, S Nagarathna, Shivaprakash M. Rudramurthy, Umabala Pamidimukkala, Anupma Jyoti Kindo

## Abstract

Primary cerebral phaeohyphomycosis (PCP) is a severe neurological infection caused by neurotropic dematiaceous fungi, affecting both immunocompetent and immunocompromised individuals. Understanding the virulence and adaptation mechanisms of these fungal pathogens is crucial for developing effective treatment strategies. This study employed Oxford Nanopore long-read sequencing to explore the genomes of *Cladophialophora bantiana*, *Fonsecaea monophora*, and *Cladosporium cladosporioides*, three key species associated with PCP. KEGG pathway analysis revealed significant enrichments in carbohydrate and amino acid metabolism, highlighting the metabolic versatility of these fungi. The analysis of transposable elements showed varying proportions of repeats, with *C. bantiana* exhibiting the highest repeat content. Additionally, the presence of diverse families of carbohydrate-active enzymes (CAZymes) emphasized their capacity for metabolizing complex carbohydrates. The analysis also identified enrichments in secondary metabolite (SM) biosynthetic gene clusters and stress adaptation pathways. All three species possess essential genes for thermal stress adaptation, such as HSP60 and HSF1, along with enzymes for detoxifying reactive oxygen species. The examination of pathogenicity-related genes uncovered a range of virulence factors, including lethal and hypervirulence genes, which raise critical concerns for human health. Functional annotations linked many of these genes to CAZymes, SMs, and stress response proteins. Furthermore, multiple efflux transporters and genes associated with antifungal resistance were identified, indicating potential adaptive mechanisms for drug resistance. This study not only advances our understanding of the genomic features of these fungi but also highlights their ecological and clinical significance, providing a foundation for future research into their pathogenicity and resistance mechanisms.

## INTRODUCTION

Primary cerebral phaeohyphomycosis (PCP) is a rare yet serious neurological infection caused by neurotropic dematiaceous fungi, characterized by darkly pigmented hyphae. Most of the causative agents belong to the order *Chaetothyriales*, with *Cladophialophora bantiana* accounting for about 50% of cases [1]. Other implicated genera include *Exophiala*, *Rhinocladiella*, *Fonsecaea*, *Exserohilum*, *Cladosporium*, *Curvularia*, and *Alternaria* [2]. While these fungi are mainly environmental saprobes causing opportunistic infections, roughly half of the affected patients are immunocompetent and have no underlying diseases, especially in cases of brain abscess [2]. Understanding the biology, lifestyle, virulence factors, and stress adaptation mechanisms of these fungi is essential for developing effective diagnostic and therapeutic strategies.

Next-generation sequencing (NGS) technologies have revolutionized microbial research, shifting the paradigm of genomics to address biological questions at a genome-wide scale. Short-read sequencing is both cost-effective and precise, supported by a variety of computational tools and pipelines [3]. However, repetitive regions in genomes can complicate reconstruction, leading to fragmented assemblies as it’s difficult to assign sequences to specific locations. In contrast, long-read sequencing can address these challenges by resolving complex genomic regions, identifying intricate structural variants, and facilitating de novo assembly [4]. Oxford Nanopore Technology (ONT) has emerged as a ground-breaking platform in mycological research, enabling real-time, long-read sequencing of fungal genomes [5]. This allows for an in-depth analysis of fungal genomic landscape, revealing insights into phylogenetic relationships, environmental adaptations, and virulence determinants [6,7]. By elucidating genomic features linked to virulence, ONT can help identify specific genes associated with pathogenicity, including those involved in melanin production and other virulence factors that enhance survival within host tissues [8]. Understanding the lifestyle and ecological roles of these fungi can provide context for their pathogenic potential, illustrating how environmental factors influence their virulence [9]. Furthermore, ONT enables researchers to investigate stress adaptation mechanisms, including resistance to oxidative stress [10] and antifungal agents [11]. This knowledge is crucial for addressing the growing challenge of antifungal resistance, as understanding how these fungi adapt to stressors can inform the development of new therapeutic approaches.

Data on the genomic landscape of neurotropic dematiaceous fungi are limited, primarily relying on traditional short-read sequencing technologies [12]. This study marks the first attempt to develop reference genome assemblies for *Cladophialophora bantiana*, *Fonsecaea monophora*, and *Cladosporium cladosporioides*, three neurotropic fungal species commonly associated with PCP. By utilizing Oxford Nanopore long-read sequencing technology, we aim to enhance our understanding of their virulence and lifestyles, offering crucial insights into their adaptation strategies that could improve diagnostic and treatment approaches for affected patients.

## Material and Methods

### Fungal strains and genomic DNA extraction

The study included three isolates of neurotropic dematiaceous fungi: *C. bantiana* CPPAIIMS, *F. monophora* CPRSRMC-02, and *C. cladosporioides* CPPNIMHANS-01, all sourced from human PCP cases. The isolates were collected from the All India Institute of Medical Sciences (Jodhpur), Sri Ramachandra Institute of Higher Education and Research (Chennai), and the National Institute of Mental Health and Neuro Sciences (Bengaluru), respectively. They were handled in a Biosafety Level 2 containment facility, in accordance with guidelines from the Centers for Disease Control and Prevention, USA. They were grown on potato dextrose agar (supplemented with 0.02% chloramphenicol) for 7 days at 25°C. The purity and viability of the cultures were ascertained by observing the colony characteristics and microscopic morphologies. The identification to the species-level was confirmed by sequencing the internal transcribed spacer (ITS) region of ribosomal DNA. The sequences were submitted to the GenBank database under the accession numbers PP462151, PP462152, and PP462153.

High-quality genomic DNA was extracted by a modification of the procedure described by Navarro-Munoz *et al* [13]. In the modified protocol, cetyltrimethylammonium bromide (CTAB) was used along with lyticase, in presence of glass beads to lyse the fungal cell wall and release the genomic DNA. Briefly, 1 cm_2_ of mycelial growth was transferred to a 2 ml Eppendorf tube containing 490 µl of 2× CTAB buffer (2% CTAB w/v, 1.4M NaCl, EDTA, 20 mM, pH 8.0, Tris.HCL, 100Mm, pH 8.0, and H_2_O), 6-10 acid-washed glass beads, 10 µl Proteinase K (10 mg/ml), and 20 U Lyticase (Sigma-Aldrich L2524-10KU) and incubated for 60 minutes at 65°C, with periodic vortexing. After incubation, 250 µl of 5% CTAB buffer was added and incubated for an additional 180 minutes at 65°C, with intermittent mixing.

Thereafter, 750 µl chloroform:isoamyl alcohol (24:1) was added and centrifuged at 14000 rpm for 10 minutes to obtain a clear supernatant containing genomic DNA. The DNA sample volume was estimated, and 0.55× volume of ice-cold 2-propanol was added and centrifuged at 12000 rpm for 10 minutes. Then, 2-propanol was carefully poured off without disturbing the pellet. The latter was subsequently washed twice with 1 ml of ice-cold 70% ethanol and centrifuged at 12000 rpm for 2 minutes. Next, the pellet was dried and re-suspended in 20-50 µl of Tris EDTA buffer (10 mM Tris, 0.1 mM EDTA, pH 8). Finally, 1 µl RNAse Cocktail Enzyme mix (AM2286; ThermoFisher Scientific, Waltham, MA, USA) per 100 µl sample was added and incubated for 1 h at 37°C.

### DNA quality control and library preparation

The concentration and purity of the extracted DNA were measured with Qubit™ dsDNA HS Assay Kit (Invitrogen) and NanoDrop 2000 spectrophotometer (ThermoFisher Scientific, Waltham, MA, USA). The integrity of the DNA was assessed on 0.35% agarose gel, alongside the Puregene NEX-GEN DNA Ladder (Genetix Biotech Asia Pvt. Ltd.) to confirm the absence of degradation. Samples were pooled and cleaned using the Zymoclean™ Large Fragment DNA Recovery Kit (Zymo Research, Los Angeles, CA, USA) following the manufacturer’s protocol. Library preparation was performed with the KAPA HyperPlus Library Preparation Kit (Kapa Biosystems, Wilmington, MA). The final libraries were quantified using a Qubit 4.0 fluorometer (ThermoFisher Scientific, Waltham, MA, USA). To determine the insert size, TapeStation 4150 (Agilent Technologies, Inc.) with high-sensitivity D1000 screentapes was used, according to the manufacturer’s instructions. Libraries were normalized to a concentration of 300 micromoles. Genomic DNA was end-polished and A-tailed using the NEBNext Ultra II End Repair/dA-Tailing Module (New England Biolabs, MA, USA). The end-prepared samples were barcoded using the Blunt/TA Ligase Master Mix (New England Biolabs, MA, USA). Sequencing adapters (SQK-NBD114.96; ONT, Oxford, UK) were ligated to the double-stranded DNA fragments using NEB Quick T4 DNA Ligase (New England Biolabs, MA, USA). After purification with AMPure XP Beads (Beckman Coulter, Inc., Indianapolis, IN, USA), the prepared library was sequenced.

### Nanopore sequencing and genome assembly

The libraries for short and long reads were quantified using a Qubit 4.0 fluorometer (ThermoFisher Scientific, Waltham, MA, USA), pooled, and sequenced with Illumina NovaSeq 6000 (Illumina), and Nanopore PromethION system (PromethION P24 and Data Acquisition Unit, ONT, Oxford, UK), respectively. For long reads, the PromethION flow cell (FLO-PRO114M R10.4.1; ONT) was selected based on the effective library concentration and data volume. A comprehensive analysis of the genomic data was performed, including genome assembly and polishing, to elucidate the genomic characteristics. Quality control of the raw sequencing data was conducted by examining read length distributions and base quality scores using NanoPlot (version 1.39.0) [14]. Adapters were trimmed with Trimmomatic (version 0.39) [15], and high-quality reads were filtered using NanoQC (version 0.9.4) for further analysis. Isolates were preliminarily identified based on the filtered high-quality reads using EukDetect (version 20220425) [16]. The filtered reads were assembled into contiguous sequences (contigs) using Canu (version 2.1.1) [17], with an input genome size of 36.72 Mbp provided to guide the assembly process. The assembled contigs were evaluated using QUAST (version 5.0.2) [18] to generate assembly statistics, including contiguity, accuracy, and completeness, followed by two rounds of polishing with Pilon (version 1.24) [19] to enhance accuracy and rectify errors. The integrity of fungal genome assembly was assessed using BUSCO (version 5.2.2) [20].

### Phylogenetic analysis

The phylogenetic analysis used internal transcribed spacer (ITS) sequences from 60 dematiaceous fungi obtained from GenBank. Sequence alignment was performed with MAFFT (version 7.490) [21]. The evolutionary history was inferred using the maximum likelihood method based on the Kimura 2-parameter model. Maximum likelihood analysis was conducted with RAxML (version 8.2.12) employing the GTR+GAMMA substitution model [22]. The bootstrap consensus tree, generated from 1000 replicates, was used to represent the evolutionary history of the analysed taxa. Initial trees for the heuristic search were obtained automatically using the Neighbor-Join and BioNJ algorithms on a matrix of pairwise distances estimated via the Maximum Composite Likelihood (MCL) approach, selecting the topology with the highest log likelihood. The phylogenetic tree was visualized using iTOL, allowing for interactive exploration and annotation of the evolutionary relationships among the fungi. Bootstrap values above 90% for Bayesian probability and above 70% for maximum likelihood were highlighted in bold to emphasize well-supported nodes.

### Gene prediction and genome functional annotation

For gene model prediction, contigs smaller than 200 bp were removed from the assembled genomes, and repeats were soft-masked using Funannotate (version 1.8.17) (https://github.com/nextgenusfs/funannotate, accessed on 16 August 2024). Gene prediction was conducted using Funannotate, which integrates ab initio gene prediction tools such as AUGUSTUS (version 3.5.0) [23], GeneMark-ES (version 4.72) [24], and SNAP (version 2006-07-28) [25]. The predicted gene models were functionally annotated by comparing them against public databases, including UniProt [26], Refseq, InterPro (version 76.0) [27], and evolutionary genealogy of genes with Non-supervised Orthologous Groups (eggNOG version 5.0.2) [28]. The BLAST results were filtered using a query coverage of > 70% and a percentage identity of >60%. The high-scoring segment pairs from the filtered BLAST results were assigned as the annotations for the gene models. Predictions for individual rRNA and tRNA were obtained using Barrnap (version 0.9) (https://github.com/tseemann/barrnap, accessed on 16 August 2024) and tRNAscan-SE (version 2.0.12) [29], respectively, while other non-coding RNAs (ncRNAs) were predicted by comparison with Rfam (version 14.0) [30]. Signal peptides were predicted using SignalP (version 6.0) [31]. The predicted gene sequences were compared with functional databases like the Kyoto Encyclopedia of Genes and Genomes (KEGG) [32] and Gene Ontology (GO) [33] for gene function annotations. Metabolic pathways were annotated using the KEGG database, with KOBAS version 3.0 [34] employed to associate KEGG orthology and pathways. The protein sequences of the predicted gene models served as input for the KEGG Mapper [35], which mapped them to KEGG orthologs using GhostKOALA [36] to reconstruct the pathways. Additionally, protein orthologs were used for de novo prediction of GO annotations via the PANNZER2 web server, employing Benjamini-Hochberg-adjusted p-values [37]. Based on the analysis from the generic GO-slim database, the GO terms were classified into three categories: biological processes (BP), molecular functions (MF), and cellular components (CC).

### Transposable elements annotation

The unique repetitive sequences in the fungal genomes were identified by homologous annotation with RepeatMasker (version 4.1.6) [38] and annotated de novo with RepeatModeler (version 2.0.5) [39] to obtain a detailed annotation of the repeats present in the query sequences, as well as a modified version in which all the annotated repeats have been masked.

### Comparative analysis of protein sequences across the three fungal species

The protein sequences of the three fungal species were retrieved and analysed for shared and unique sequences using OrthoVenn3 (2022) [40], a web-based tool for identifying and annotating orthologous clusters across multiple species based on pairwise sequence similarities. To filter out non-significant similarities, an e-value cutoff of 1e−5 was applied, and the remaining similar sequences were organized into orthologous clusters with an inflation value of 1.5.

### Carbohydrate-active enzymes annotation

CAZy is a database of Carbohydrate-Active enZymes (CAZymes), containing a sequence-based classification and associated information about enzymes involved in the synthesis, metabolism, and recognition of complex carbohydrates [41]. In this study, the genes encoding carbohydrate-active enzymes (CAZymes) were predicted with HMMER (version 3.4) [42] against profiles from the database of automated Carbohydrate-active enzyme ANnotation (dbCAN) version 12.0 [43], which is based on the CAZy database. The CAZymes were classified according to the amino acid sequence similarities: glycosyltransferases (GTs), glycoside hydrolases (GHs), polysaccharide lyases (PLs), carbohydrate esterases (CEs), carbohydrate-binding modules (CBMs), and auxiliary activity redox enzymes (AAs). The data for *C. bantiana*, *F. monophora* were obtained directly from the CAZy database, while for *C. cladosporioides*, data from its closest relative, *C. sphaerospermum* UM 843, were utilized due to the unavailability of species-specific data. Filtering criteria were applied based on sequence identity percentages, with thresholds set at 90% for *C. bantiana* and *F. monophora*, and 80% for *C. cladosporioides*, considering its close relationship with the selected species. This approach ensured a comparative analysis of CAZymes, providing insights into their functional roles and evolutionary relationships in carbohydrate metabolism.

### Extracellular peptidases annotation

The extracellular peptidase-related genes, such as serine peptidase (SP), aspartic peptidase (AP), metallopeptidase (MP), cysteine peptidase (CP), threonine peptidase (TP), and glutamic peptidase (GP) were predicted through a BLAST search against the MEROPS database (version 12.5) [44]. This analysis aimed to identify the peptidase sequences in three species with an identity percentage above 90% and an e-value of less than 1e-5. To ensure relevance, inhibitor sequences were excluded from the analysis.

### Cytochrome P450 annotation

Cytochrome P450 (CYP) enzymes are fundamental to fungal stress adaptation and detoxification of exogenous toxic compounds (e.g. environmental pollutants, toxins, and xenobiotics). They are also involved in various stages of fungal development, virulence and pathogenesis. In this study, BLASTP (version 2.12.0) was used for CYP annotation based on the Fungal Cytochrome P450 Database (FCPD) [45], a repository exclusively meant for cataloguing and studying CYP enzymes in fungi.

### Transmembrane proteins annotation

The protein sequences of all the predicted genes were analysed by SignalP (version 6.0) [31], and those containing signal peptides were checked for transmembrane helices using TMHMM (version 2.0) [46]. The output files were filtered based on a significance threshold (p ≤0.05), followed by another filtration step, retaining entries with a predicted number of transmembrane helices ≥1.

### Secondary metabolites annotation

For analysis of secondary metabolite (SM) biosynthetic gene clusters, antiSMASH (version 3.0) [47] was used. The data for *C. bantiana* was obtained directly from the database, while for *F. monophora* and *C. cladosporioides*, BLAST analysis was performed against *F. pedrosoi* and *Aureobasidium namibiae* due to the unavailability of species-specific data. Sequences with identity percentages above 90% were retained for *C. bantiana* and *F. pedrosoi*, while for *C. cladosporioides*, sequences with identity percentages exceeding 60% were selected.

### Iron uptake and homeostasis annotation

The key elements involved in fungal iron uptake and homeostasis include L-ornithine-N5-monooxygenase, ferricrocin, siderophore transporters, the Sid C, Sid D, Sid F, and Sid G genes, a GATA-like transcription factor, and the ferroxidase Fet3 in conjunction with the iron permease Ftr1. The sequences related to these components were identified and analysed using BLASTP (version 2.12.0). The results were filtered with an identity percentage above 60% to ensure relevance.

### Stress adaptation pathways annotation

To delineate the stress biology, the potential protein-coding genes related to fungal stress adaptation, such as heat shock proteins (HSPs), superoxide dismutase (SOD), catalase-peroxidase, thioredoxin reductase, monothiol glutaredoxin, glutathione peroxidase, methionine sulfoxide reductase, alpha-trehalose-6-phosphate synthase and mitogen-activated protein kinases (MAPK) were analysed by BLASTP (version 2.12.0), using the fungal stress response database (FSRD) [48].

### Virulence factors annotation

The Database of Fungal Virulence Factors (DFVF) is a valuable resource that catalogs genes linked to fungal virulence [49]. This database was accessed to gather all genes identified as potential virulence factors. Additionally, the Pathogen-Host Interactions Database (PHI-base version 4.12) [50], which is a comprehensive resource documenting experimentally verified pathogenicity, virulence, and effector genes, was also searched. The complete dataset from PHI-base was downloaded to aid in identifying hypervirulence and lethal genes in the species of interest. The gene IDs from the filtered results were used to extract corresponding sequences representing fungal genes potentially involved in human infections for subsequent analyses. To assess the presence and similarity of these human-associated fungal genes across three species, the extracted sequences were analysed using BLAST search against the genomic data of the three species. The results were filtered to retain only hits with an identity percentage above 70%, ensuring that only highly similar sequences were considered for further analysis. A keyword search was performed on the filtered BLAST results to classify genes into hypervirulent and lethal categories.

### Antifungal resistance genes annotation

To identify the potential genes associated with antifungal resistance, the protein-coding sequences of the three species were aligned using BLASTP (version 2.12.0) against the Mycology Antifungal Resistance Database (MARDy) [51], specifically targeting the genes ERG11/CYP51a, FKS1, FKS2, and FUR1. In addition, the sequences for efflux transporters associated with multidrug resistance in fungi, such as ATP-binding cassette (ABC) transporters (MDR1, CDR1, and CDR2) and Major Facilitator Superfamily (MFS) transporters (DHA1 and DHA2) were also analysed. The results were filtered with identity percentages exceeding 50%.

## RESULTS

### Sequence assembly and genomic features

The genomes of *C. bantiana*, *F. monophora*, and *C. cladosporioides* were assembled using the genomic DNA sequences generated by nanopore long-read technology, which constituted 9.76 Gb, 5.12 Gb and 6.79 Gb of data, respectively. The sequenced reads were analysed using K-mer to estimate genome size and heterozygosity. The genome sizes obtained for *C. bantiana*, *F. monophora*, and *C. cladosporioides* were 39.9 Mb (39,883,971 bp), 35.9Mb (35,921,295 bp), and 31.5 Mb (31,487,539 bp), comprising of 30, 15, and 22 contigs, respectively. The N50 contig lengths were approximately 4.8 Mb, 5.2 Mb, and 1.87 Mb, and consisted of 4, 3, and 7 scaffolds with an overall GC content of 51.19%, 52.16%, and 52.9%, respectively. BUSCO (version 4.1.4) analysis of *C. bantiana*, *F. monophora*, and *C. cladosporioides* genome assemblies based on the representation of core eukaryotic genes (Fungi Odb10 datasets) indicated that the genomes were highly complete, with 99.1%, 98.9%, and 99.3% of these genes present in all assemblies, respectively. The genome assembly results are summarized in Table 1.

**Table 1.** Sequencing and assembly features.

| Details | *C. bantiana* | *F. monophora* | *C. cladosporioides* |
| --- | --- | --- | --- |
| Sequencing depth | 90× | 90× | 90× |
| Total length of sequences (bp) | 39,883,971 | 35,921,295 | 31,487,539 |
| Number of reads | 1,874,839 | 2,925,125 | 8,542,468 |
| Mean read length (bp) | 5206.9 | 1751.2 | 795.5 |
| Number of contigs ($\geq 1000$ bp) | 30 | 15 | 22 |
| Number of contigs ($\geq 5000$ bp) | 28 | 14 | 18 |
| Number of contigs ($\geq 10000$ bp) | 23 | 14 | 17 |
| Number of contigs ($\geq 25000$ bp) | 19 | 12 | 17 |
| Number of contigs ($\geq 50000$ bp) | 15 | 09 | 17 |
| Longest contig (bp) | 7,105,748 | 8,543,123 | 3,517,742 |
| GC content (%) | 51.19 | 52.16 | 52.90 |
| Contig N50 (bp) | 4810828 | 5200119 | 1871712 |
| Contig L50 | 4 | 3 | 7 |
| Contig N90 (bp) | 3999628 | 4324844 | 1468201 |
| Contig L90 | 7 | 6 | 15 |
| auN | 4931794.0 | 5670806.3 | 2051600.5 |
| Complete and single-copy BUSCO (%) | 99.1 | 98.9 | 99.3 |
| Complete and duplicated BUSCO (%) | 0 | 0.1 | 0.3 |
| Fragmented BUSCO (%) | 0.8 | 0.5 | 0 |
| Missing BUSCO (%) | 0.1 | 0.5 | 0.4 |

BUSCO, Benchmarking Universal Single-Copy Orthologs

The genome data analysed using AUGUSTUS v3.5.0 gene prediction tool identified a total of 12,680, 11,736, and 10,773 putative coding sequences (CDS) in the genomes of *C. bantiana*, *F. monophora*, and *C. cladosporioides*, respectively. The average sizes of protein coding genes were 1,407 bp, 1,504 bp, and 1,119 bp, respectively. On an average, there were 2.3, 2.4, and 2.2 exons per gene, with mean exon sizes of 625 bp, 620 bp, and 671 bp, respectively. The average numbers of introns per gene were 1.3, 1.4, and 1.2, respectively. A total of 148, 65, and 301 non-coding RNAs were predicted in the respective genomes, with tRNA having the highest copy number and consisting mainly of 5S rRNA, 5.8S rRNA, 18S rRNA, and 28S rRNA. There were 27, 2 and 3 ITS regions in the respective genomes (Table 2).

**Table 2.**
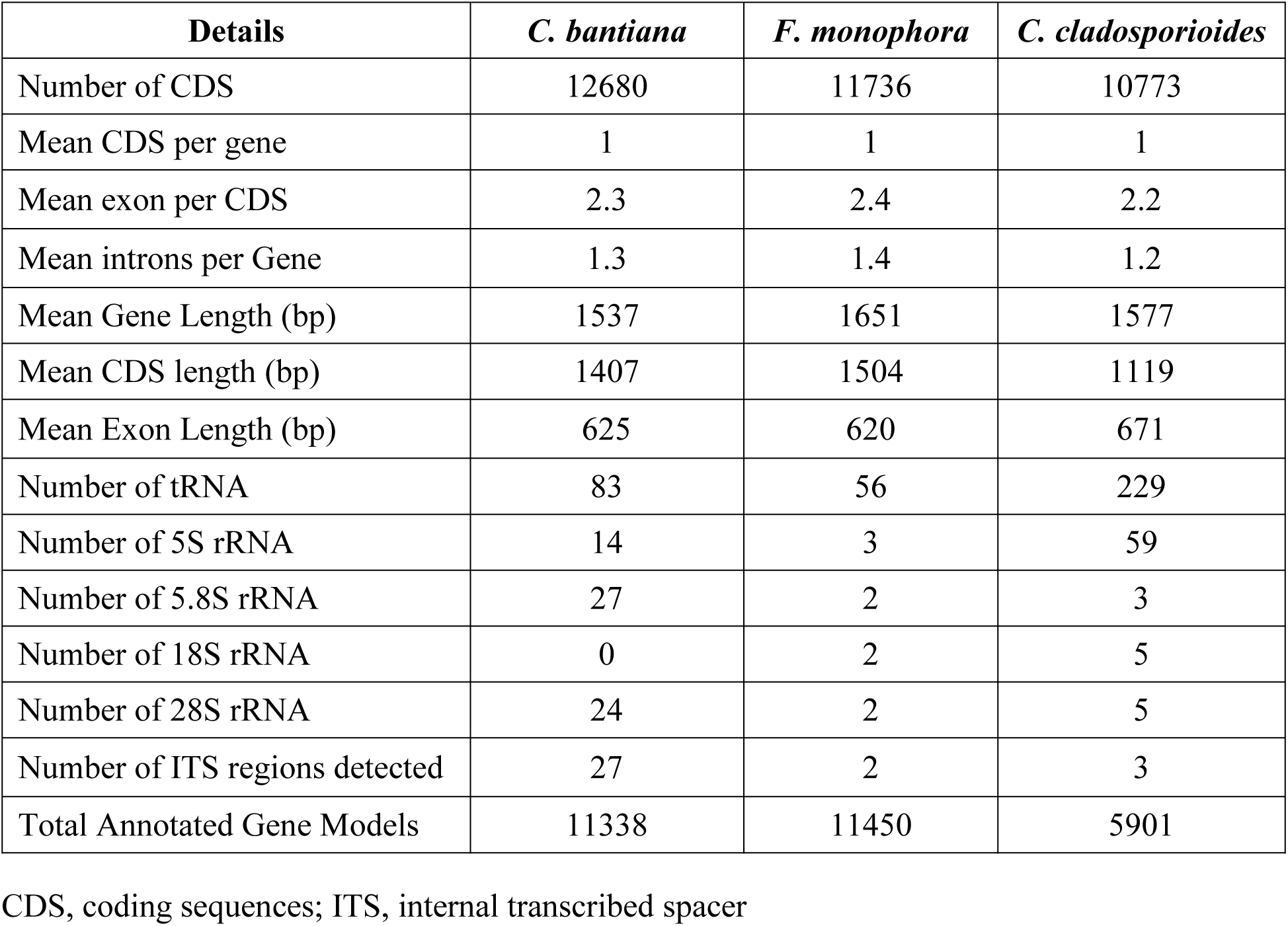
Gene prediction summary.

| Details | *C. bantiana* | *F. monophora* | *C. cladosporioides* |
| --- | --- | --- | --- |
| Number of CDS | 12680 | 11736 | 10773 |
| Mean CDS per gene | 1 | 1 | 1 |
| Mean exon per CDS | 2.3 | 2.4 | 2.2 |
| Mean introns per Gene | 1.3 | 1.4 | 1.2 |
| Mean Gene Length (bp) | 1537 | 1651 | 1577 |
| Mean CDS length (bp) | 1407 | 1504 | 1119 |
| Mean Exon Length (bp) | 625 | 620 | 671 |
| Number of tRNA | 83 | 56 | 229 |
| Number of 5S rRNA | 14 | 3 | 59 |
| Number of 5.8S rRNA | 27 | 2 | 3 |
| Number of 18S rRNA | 0 | 2 | 5 |
| Number of 28S rRNA | 24 | 2 | 5 |
| Number of ITS regions detected | 27 | 2 | 3 |
| Total Annotated Gene Models | 11338 | 11450 | 5901 |

CDS, coding sequences; ITS, internal transcribed spacer

### Phylogeny

Multilocus phylogenetic analysis of three species using the ITS region of rDNA revealed that *C. bantiana* CPPAIIMS (PP462151) and F. monophora CPRSRMC-02 (PP462152) clustered within members of the order *Chaetothyriales*, while *C. cladosporioides* CPPNIMHANS-01 (PP462153) grouped among members of *Cladosporiales* in a paraphyletic position. An ITS-based phylogenetic tree was constructed, including representatives from the orders *Umbilicariales*, *Erysiphales*, *Pleosporales*, *Helotiales*, *Venturiales*, *Sordariales*, Mycosphaerellales, and *Cladosporiales*, with species from *Chaetothyriales* serving as the focus (Fig. 1). The family comprising the genera *Camptophora*, *Arthrocladium*, *Capronia*, and *Knufia* was clearly distinguished from the other genera within *Chaetothyriales*. In the ITS tree, *C. bantiana* CPPAIIMS formed a separate, well-supported cluster (100% bootstrap) with two other strains, while *F. monophora* CPRSRMC-02 clustered with three additional strains in a distinct group, also with 100% bootstrap support. The strain *C. cladosporioides* CPPNIMHANS-01 clustered with seven other strains from *Cladosporiales* with 99% bootstrap support. No distinct ecological trends were observed between the strains. *Placocarpus schaereri* (EU006532), a member of the order *Verrucariales*, was used as an outgroup.

**Figure 1.**
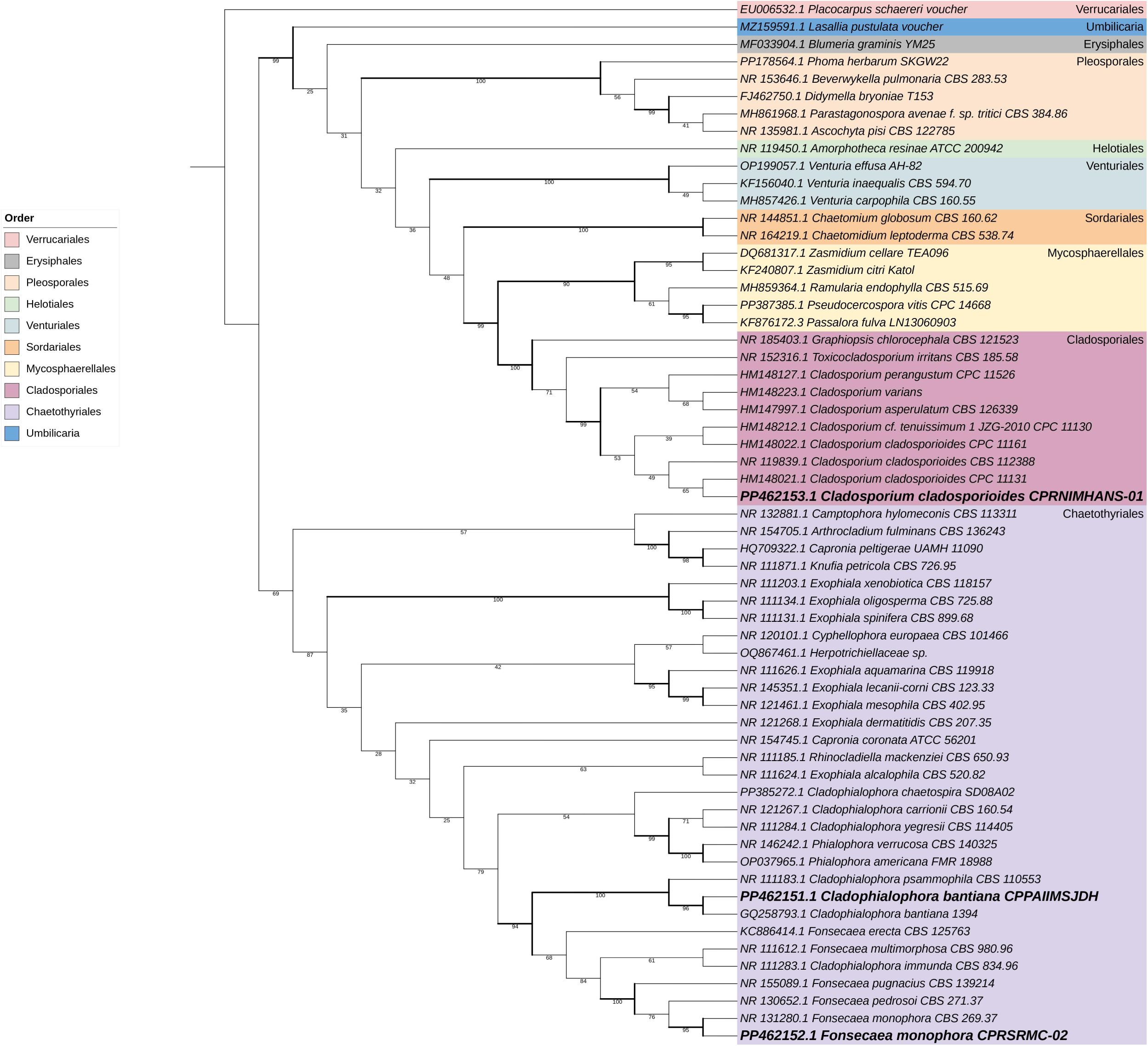
Phylogenetic analysis of *C. bantiana* CPPAIIMS, *F. monophora* CPRSRMC-02, and *C. cladosporioides* CPPNIMHANS-01. The Maximum likelihood tree, based on 60 ITS sequences, was determined using RAxML v. 8.2.12 with Kimura 2-parameter model. The bootstrap consensus tree, generated from 1000 replicates, was used to represent the evolutionary history of the analyzed taxa. The tree was rooted to *Placocarpus schaereri* (EU006532).

### KEGG functional annotation of *C. bantiana*, *F. monophora*, and *C. cladosporioides* genes

We performed KEGG pathway analysis to gain more insights into gene functions across the three species (Table S1). A total of 2,913 predicted proteins were mapped to their orthologous genes in the KEGG metabolic pathways for *C. bantiana* (Fig. 2A), 3,124 for *F. monophora* (Fig. 2B), and 2,953 for *C. cladosporioides* (Fig. 2C). The pathways with maximum enrichment included carbohydrate metabolism, amino acid metabolism, lipid metabolism, secondary metabolites biosynthesis, microbial metabolism in diverse environments, xenobiotics biodegradation, and carbon metabolism. In the “human disease” category (DFVF-HUM), we identified 79 genes associated with 22 KEGG pathways related to human infectious diseases in *C. bantiana* (Fig. 3A and D), 74 in *F. monophora* (Fig. 3B and E), and 75 in *C. cladosporioides* (Fig. 3C and F). The pathway with the highest gene enrichment was KO:05165 (Human papillomavirus infection), followed by KO:05110 (*Vibrio cholerae* infection) and KO:05120 (*Helicobacter pylori* infection) (Table S2).

**Figure 2.**
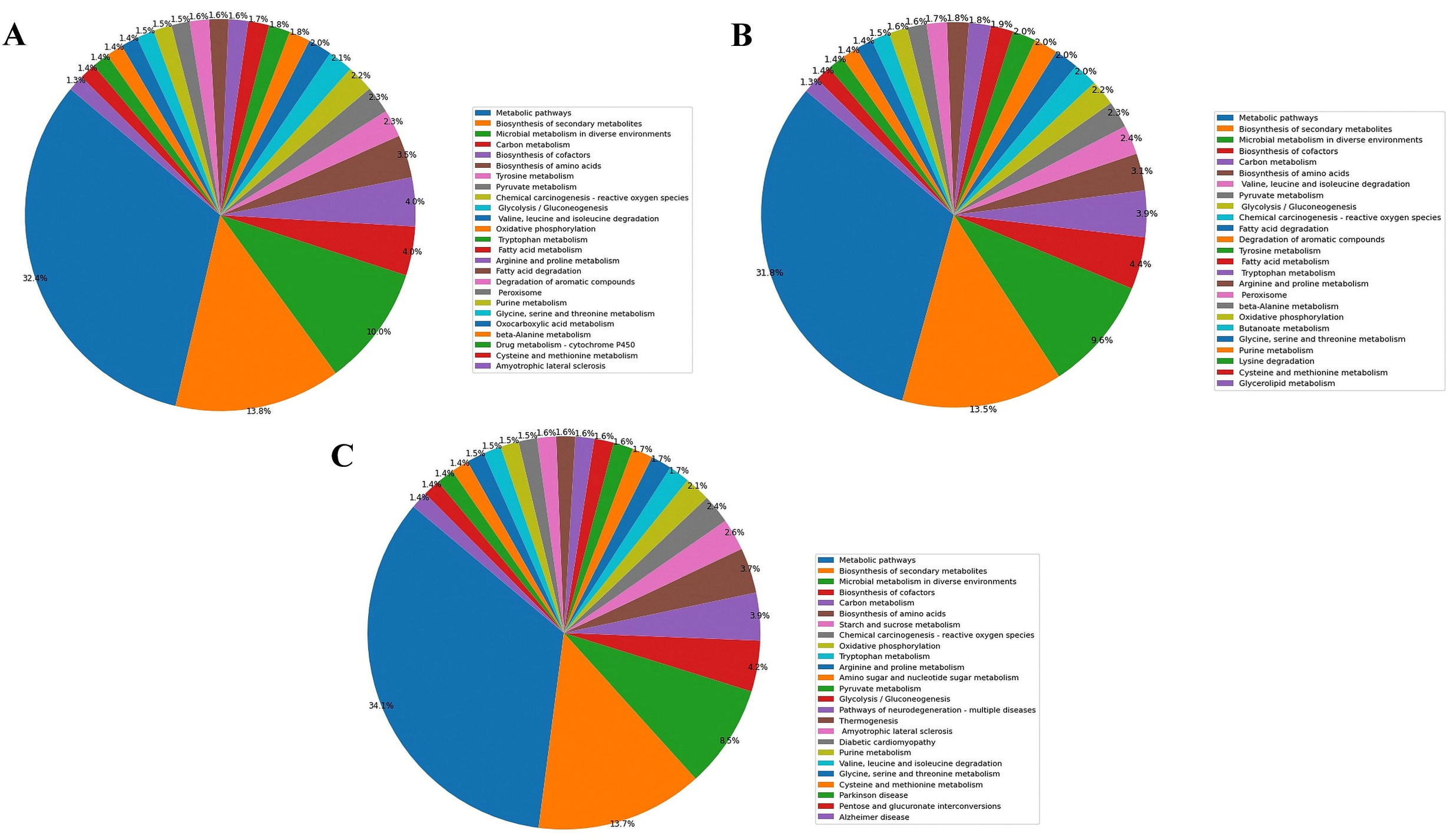
KEGG pathway analysis showing the top 25 significantly enriched pathways in *C. bantiana* (A), *F. monophora* (B), and *C. cladosporioides* (C). These pathways were identified by annotating the protein-coding sequences against the KEGG database.

**Figure 3.**
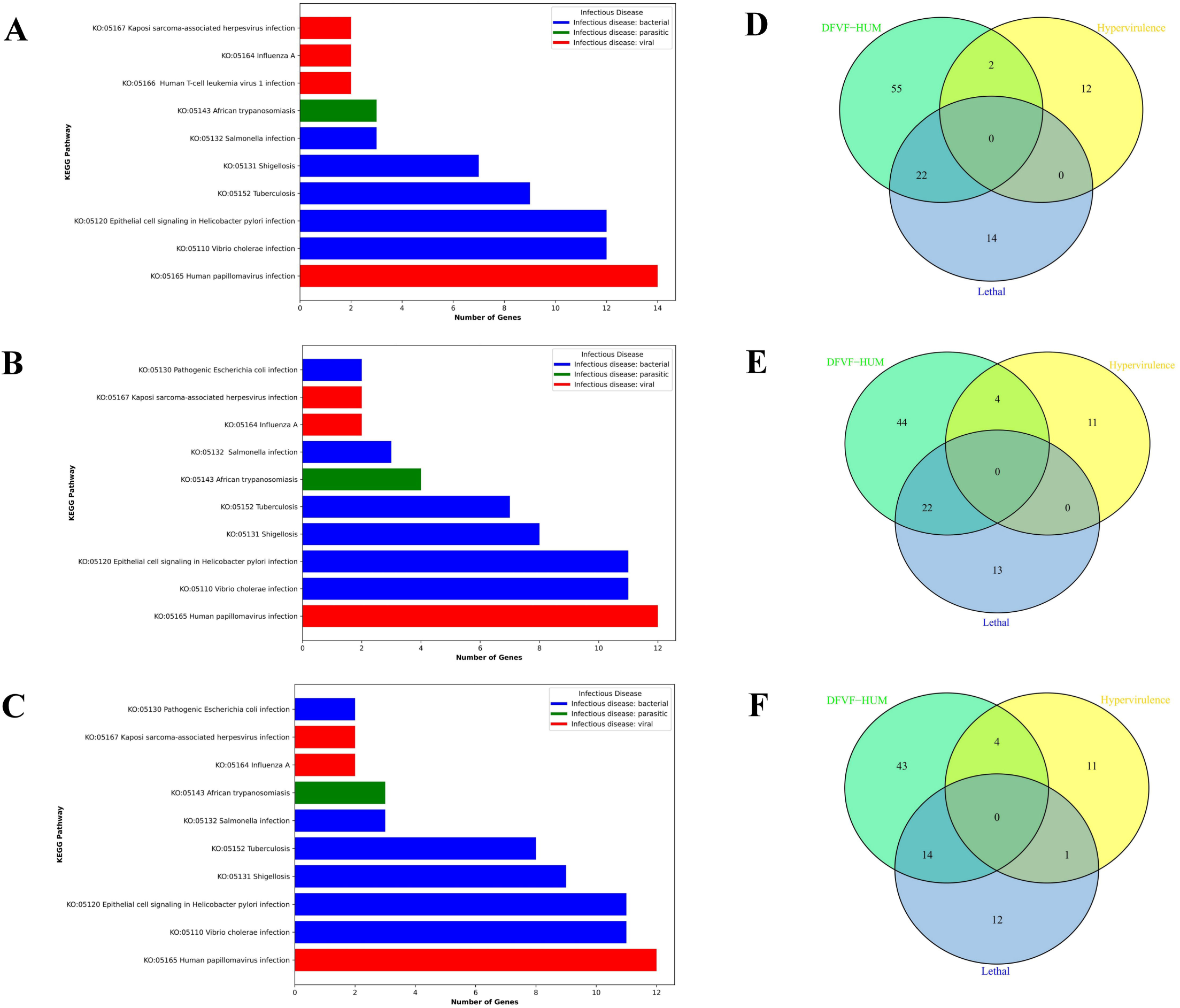
Analysis of human infectious disease KEGG pathways in *C. bantiana* (A), *F. monophora* (B), and *C. cladosporioides* (C). Venn diagrams illustrate the genes encoding DFVF-HUM, lethal virulence factors, and hypervirulence factors in *C. bantiana* (D), *F. monophora* (E), and *C. cladosporioides* (F). The genes were categorized as hypervirulent and lethal using the PHI-base.

### GO functional annotation of C. bantiana, F. monophora, and C. cladosporioides genes

In *C. bantiana*, analysis of GO terms from the GO-Slim database revealed the highest enrichment in the “molecular function” category (4,820 genes), followed by “biological process” (3,301 genes), and “cellular component” (2,318 genes). As illustrated in Fig. S1A and S2A, the primary enrichments include “membrane” (1,610), “nucleus” (561), “ATP binding” (519), “transmembrane transport” (496), and “DNA binding” (404).

In *F. monophora*, a total of 5,471 genes were classified under “molecular function,” with 3,632 in “cellular component” and 2,731 in “biological process.” Figures S1B and S2B demonstrate significant enrichments in “membrane” (1,758), “nucleus” (665), “transmembrane transport” (602), “ATP binding” (561), “zinc ion binding” (513), and “DNA binding” (500).

For *C. cladosporioides*, 1,846 genes were associated with “molecular function,” 1,470 with “cellular component” and 991 with “biological process.” As shown in Figures S1C and S2C, the key enrichments were in “membrane” (531), “ATP binding” (344), “nucleus” (201), “transmembrane transport” (196), and “metal ion binding” (190). Additional details on KEGG pathway enrichment and GO functional annotations for the predicted genes can be found in Table S3. The genomic characteristics of the three species are illustrated in circular genome maps (Fig. 4).

**Figure 4.**
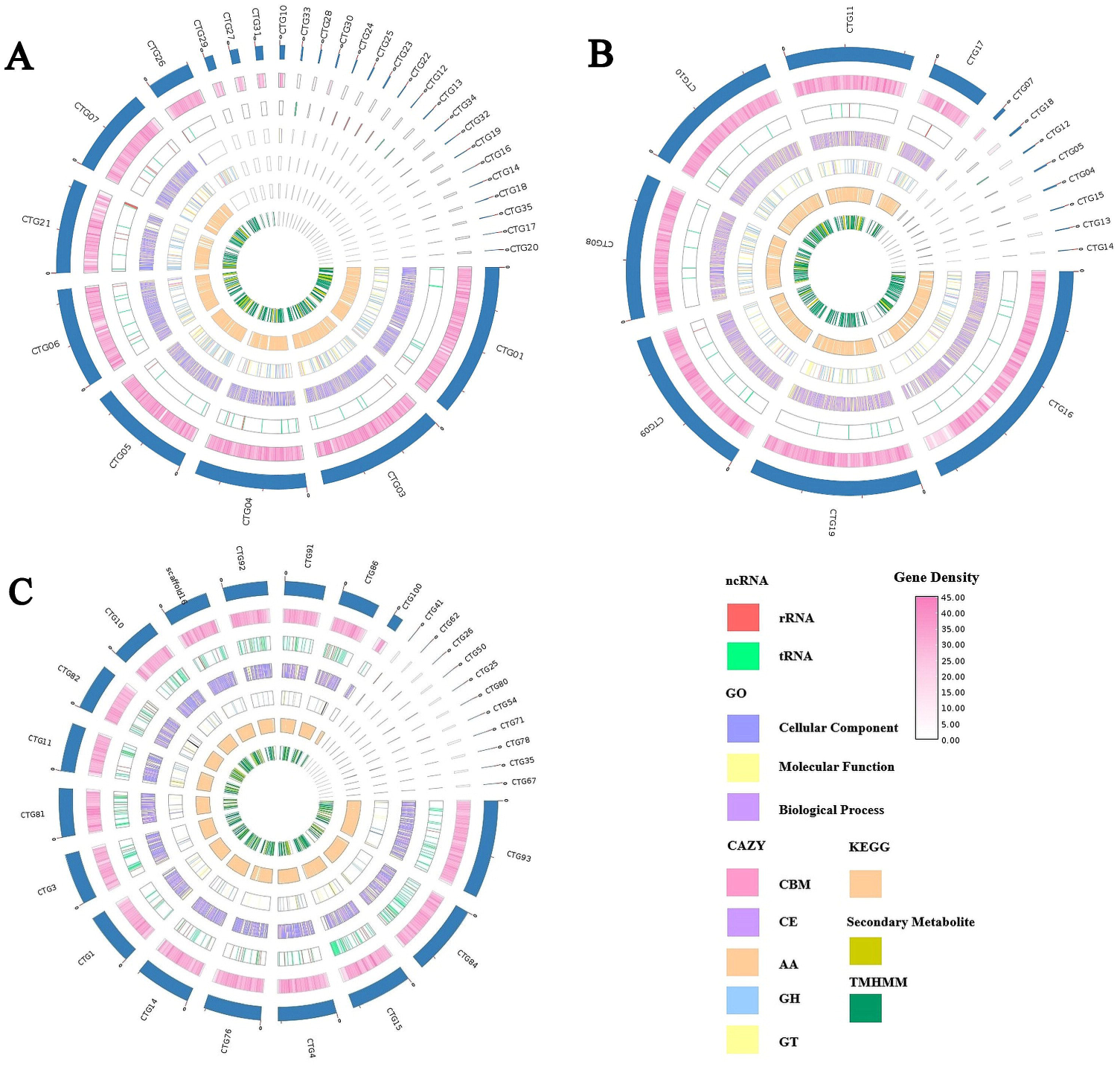
Circular genome maps of *C. bantiana* (A), *F. monophora* (B), and *C. cladosporioides* (C). From outside to the center: Circle 1 shows the contig keyboard; Circle 2 illustrates gene density; Circle 3 represents non-coding RNA (ncRNA); Circles 4-7 display predicted protein-coding genes based on the GO, CAZy, KEGG, and TMHMM databases, respectively. The distinct colors indicate different functional classifications.

### Comparative analysis of protein sequences

Using OrthoVenn3, we identified 5,660 shared protein sequences among the three species. *C. bantiana* had 3,908 unique sequences, *F. monophora* had 3,749, and *C. cladosporioides* had 745 (Fig. S3A). Additionally, we found 9,568 orthologous clusters (OCs) and 2,473 singletons in *C. bantiana*, 9,409 OCs and 1,790 singletons in *F. monophora*, and 6,405 OCs and 1,790 singletons in *C. cladosporioides* (Fig. S3B).

### Predicted transposable elements

In the genome of *C. bantiana*, 3,051,989 bp of repeat sequences were identified, comprising 7.66% of the total genome, with interspersed repeats at 5.08% and tandem repeats at 0.63%. Specifically, 696 Class I retroelements and 148 Class II DNA transposons represented 4.54% and 0.54% of the genomic length, respectively, with Gypsy/DIRS1 being the predominant retroelement at 4.18%. For DNA transposons, Helitron and Tc1-Mariner accounted for 0.32% and 0.22%, respectively. Tandem repeats were primarily composed of simple and low-complexity repeats. In the *F. monophora* genome, 9,987 TEs accounted for 3.3% of the genome, with interspersed repeats at 1.36% and tandem repeats at 0.98%; Class I and Class II TEs numbered 368 and 27, representing 1.28% and 0.08%, respectively, featuring only Gypsy/DIRS1 among retrotransposons and Tc1-Mariner among DNA transposons. Tandem repeats included simple repeats and low-complexity repeats at 0.88% and 0.1%, respectively. The *C. cladosporioides* genome revealed 6,068 TEs, making up 1.49% of the genome, with interspersed repeats at 0.35% and tandem repeats at 0.7%. It contained 45 Class I retroelements and 66 Class II TEs, corresponding to 0.15% and 0.2%, with no LTR retrotransposons identified, while Tc1-Mariner and Helitron comprised 0.19% and 0.01% of the genomic length. Simple repeats and low-complexity tandem repeats accounted for 0.64% and 0.06%, respectively. The analysis of TEs is summarized in Table 3.

**Table 3.** Statistical analysis of repeat sequences.

| Categories | C. bantiana |  |  | F. monophora |  |  | C. cladosporioides |  |  |
| --- | --- | --- | --- | --- | --- | --- | --- | --- | --- |
|  | UE | CL (bp) | GA (%) | UE | CL (bp) | GA (%) | UE | CL (bp) | GA (%) |
| **Retrotransposons (Class I)** |  |  |  |  |  |  |  |  |  |
| **1) LTR** | 623 | 1,681,233 | 4.22 | 368 | 460,103 | 1.28 | 0 | - | 0 |
| a) Gypsy/DIRS1 | 581 | 1,668,732 | 4.18 | 368 | 460,103 | 1.28 | 0 | - | 0 |
| b) Retroviral | 42 | 12,501 | 0.03 | 0 | - | 0 | 0 | - | 0 |
| **2) SINEs** | 0 | - | 0 | 0 | - | 0 | 32 | 4,942 | 0.02 |
| **3) LINEs** | 73 | 128,377 | 0.32 | 0 | - | 0 | 13 | 41,177 | 0.13 |
| **Total retroelements** | 696 | 1,809,610 | 4.54 | 368 | 460,103 | 1.28 | 45 | 46,119 | 0.15 |
| **DNA transposons (Class II)** |  |  |  |  |  |  |  |  |  |
| Tc1-IS630-Pogo | 103 | 87,924 | 0.22 | 27 | 29,360 | 0.08 | 49 | 61,045 | 0.19 |
| Rolling-circles | 45 | 126,218 | 0.32 | 0 | - | 0 | 17 | 1,600 | 0.01 |
| **Other genomic repeat elements** |  |  |  |  |  |  |  |  |  |
| Unclassified | 1,849 | 770,248 | 1.93 | 681 | 343,798 | 0.96 | 599 | 125,012 | 0.4 |
| Small RNA | 43 | 9,235 | 0.02 | 0 | - | 0 | 105 | 11,279 | 0.04 |
| Simple repeats | 5,441 | 218,546 | 0.55 | 8,155 | 317,720 | 0.88 | 4,822 | 201,047 | 0.64 |
| Low complexity | 633 | 30,208 | 0.08 | 756 | 34,636 | 0.10 | 431 | 20,410 | 0.06 |
| **Total TEs** | 8,810 | 3,051,989 | 7.66 | 9,987 | 1,185,617 | 3.3 | 6,068 | 466,512 | 1.49 |

UE, Unique Elements; CL, Cumulative Length; GA, Genome Assembly; SINE, Short Interspersed Nuclear Element; LINE, Long Interspersed Nuclear Element; LTR, Long Terminal Repeat; DIRS1, Dictyostelium Intermediate Repeat Sequence

### Carbohydrate active enzymes (CAZymes)

The CAZy annotation pipeline revealed 298 putative CAZyme genes in the *C. bantiana* genome, classified into 102 families. This includes 141 GHs, 91 GTs, 49 AAs, and 6 CEs. Additionally, 11 genes from 6 families encoding CBMs were identified (Table S4 and Fig. S4A). In the *F. monophora* genome, 275 putative CAZyme genes were identified, also across 102 families, including 123 GHs, 89 GTs, 43 AAs, and 8 CEs. Furthermore, 12 CBM genes from 8 families were detected. Neither *C. bantiana* nor *F. monophora* genomes contained any PL genes. The *C. cladosporioides* genome contained 114 putative CAZyme genes across 60 families, comprising 53 GHs, 26 GTs, 16 AAs, 7 PLs, and 5 CEs, along with 7 genes from 4 families encoding CBMs. The CAZyme annotation distributions of three species are shown in Table S5.

### Extracellular peptidases, Cytochrome P450, and Transmembrane proteins

Using the MEROPS database, 193 genes encoding extracellular peptidases were identified in *C. bantiana*, which include 60 SPs, 15 APs, 51 MPs, 47 CPs, 19 TPs, and one GP. In *F. monophora*, we found 192 genes encoding extracellular peptidases, comprising 62 SPs, 15 APs, 52 MPs, 44 CPs, 18 TPs, and one GP. For *C. cladosporioides*, 32 genes encoding extracellular peptidases were identified, consisting of eight SPs, one AP, nine MPs, three CPs, and 11 TPs, with no genes encoding GP (Table S6).

We also identified 89 unique hypothetical proteins containing the CYP domain in *C. bantiana*, 36 in *F. monophora*, and 4 in *C. cladosporioides* through annotation using the FCPD (Table S7). Additionally, we found 299 genes encoding 1,169 transmembrane proteins (TMPs) in *C. bantiana*, 276 genes for 1,278 TMPs in *F. monophora*, and 287 genes for 1,295 TMPs in *C. cladosporioides* (Table S8).

### Secondary metabolite biosynthetic gene clusters

The analysis of SM biosynthetic gene clusters using the antiSMASH revealed 338 gene clusters in the *C. bantiana* genome, categorized into nine groups: 99 nonribosomal peptide synthetase clusters (NRPS)-like clusters, 94 fungal-ribosomally synthesized and post-translationally modified peptides (RiPP)-like clusters, 29 NRPS-fungal RiPP-like clusters, 29 terpene clusters, 26 type I polyketide synthase (T1PKS) clusters, 22 betalactone-fungal RiPP-like clusters, 15 terpene-T1PKS clusters, 14 type III polyketide synthase (T3PKS) clusters, and 10 nonribosomal peptide (NRP)-metallophore-NRPS clusters.

In the *F. monophora* genome, 388 gene clusters related to SMs were identified across ten categories: 127 fungal-RiPP-like clusters, 98 NRPS-like clusters, 41 TIPKS clusters, 38 terpene clusters, 21 betalactone-fungal RiPP-like clusters, 17 NRPS clusters, 16 isocyanide-NRP clusters, 14 T3PKS clusters, 9 NRP-metallophore-NRPS clusters, and 8 betalactone clusters (Table 4 and Fig. S4B).

**Table 4.** Analysis of secondary metabolite biosynthetic gene clusters.

| Category | Number of genes |  |  |
| --- | --- | --- | --- |
|  | C. bantiana | F. monophora | C. cladosporioides |
| NRPS-like | 99 | 98 | 7 |
| Fungal-RiPP-like | 94 | 127 | 5 |
| Terpene | 29 | 38 | 1 |
| NRPS,fungal-RiPP-like | 29 | 0 | 0 |
| T1PKS | 26 | 41 | 2 |
| Betalactone,Fungal-RiPP-like | 22 | 21 | 0 |
| T3PKS | 14 | 13 | 1 |
| NRP-metallophore-NRPS | 10 | 9 | 0 |
| Terpene-T1PKS | 15 | 0 | 0 |
| Betalactone | 0 | 8 | 0 |
| Isocyanide-NRP | 0 | 16 | 0 |
| NRPS | 0 | 17 | 1 |
| NRPS-like-T1PKS | 0 | 0 | 3 |
| NAPAA | 0 | 0 | 1 |

NRP, Nonribosomal peptide; NRPS, Nonribosomal peptide synthetase; RiPP, Ribosomally synthesized and post-translationally modified peptides; T1PKS, Type I polyketide synthase; T3PKS, Type III polyketide synthase; NAPAA, non-alpha poly-amino acid

The *C. cladosporioides* genome contained 21 SM biosynthetic gene clusters, with the largest number being NRPS-like clusters (7), followed by fungal-RiPP-like clusters (5), T1PKS-NRPS-like clusters (3), T1PKS clusters (2), and one each of terpene, T3PKS, NRPS, and non-alpha poly-amino acid (NAPAA) clusters. Further details on the annotations of SM biosynthetic gene clusters can be found in Table S9.

### Iron uptake and Homeostasis

We identified 30 genes related to iron uptake and homeostasis in *C. bantiana*. The siderophore iron transporter (SIT) exhibited the highest number of gene clusters (8), followed by cupin domain-containing proteins (7), metalloreductases (4), pH response regulators (PalF/RIM8, PalA/RIM20, and PalH/RIM21) and vacuolar transporters (3 each), mitogen-activated protein kinases (MAPK) (2), and arginase and L-ornithine-N_5_-monooxygenase (1 each). Notably, a gene encoding a sterol regulatory element binding protein (NMHCB_008088) was identified in the *C. bantiana* genome, which was absent in the other two species.

In *F. monophora*, eight genes were linked to iron uptake and homeostasis, primarily involving SIT, MAPK, vacuolar transporter, arginase, and L-ornithine-N_5_-monooxygenase.

In *C. cladosporioides*, eight genes associated with iron uptake and homeostasis were identified, involving key pathways like MAPK, pH response regulators (PalA/RIM20 and PalH/RIM21), and vacuolar transporters. Unlike *C. bantiana* and *F. monophora*, *C. cladosporioides* lacks genes for arginase and L-ornithine-N5-monooxygenase. Notably, a unique high-affinity iron permease (NMHCC_006316) was identified. A summary of the genes and pathways involved in iron uptake and homeostasis is provided in Table S10.

### Stress adaptation pathways

*C. bantiana*, *F. monophora*, and *C. cladosporioides* were predicted to contain HSP60 and heat shock transcription factor 1 (HSF1) in their genomes, both of which are essential for cellular adaptation to thermal stress. Additionally, all three species were found to possess several genes encoding proteins that play crucial roles in detoxifying reactive oxygen species (ROS) produced by human phagocytes during infection, including SOD, glutathione peroxidase, catalases, components of the thioredoxin and glutaredoxin systems, and MAPKs. The genes for nitric oxide dioxygenase and trehalose-6-phosphate synthase were present in *C. bantiana* and *F. monophora*, but were absent in *C. cladosporioides*. Interestingly, *C. cladosporioides* also contained two unique genes: a zinc cluster transcription factor (ZCTF) (NMHCC_001798) and a methionine sulfoxide reductase (NMHCC_009623) (Table 5).

**Table 5.** Analysis of stress adaptation pathways.

| Stress response pathways | Number of genes |  |  |
| --- | --- | --- | --- |
|  | **C.**<br>**bantiana** | **F. monophora** | **C. cladosporioides** |
| Heat shock protein 60 | 1 | 1 | 1 |
| Heat shock transcription factor | 1 | 1 | 1 |
| Nitric oxide dioxygenase | 1 | 2 | 0 |
| pH response regulator protein | 3 | 0 | 2 |
| Superoxide dismutase 2 | 6 | 3 | 3 |
| Monothiol glutaredoxin 3 | 2 | 2 | 1 |
| Glutathione peroxidase | 1 | 1 | 1 |
| Catalase | 9 | 7 | 1 |
| Mitogen-activated protein kinase | 7 | 3 | 8 |
| Trehalose-6-phosphate synthase | 4 | 2 | 0 |
| Methionine sulfoxide reductase | 0 | 0 | 1 |
| Zinc cluster transcription factor | 0 | 0 | 1 |

### Pathogenicity-related genes and virulence factors

A comprehensive whole-proteome BLAST search against the PHI-base identified several pathogenicity-associated genes categorized into six main groups (Table S11). In *C. bantiana*, 68 virulence-related genes were identified, with 66 linked to reduced virulence and two (NMHCB_003848 and NMHCB_009577) associated with hypervirulence. Additionally, there were 11 genes classified as “lethal” and two involved in “resistance to chemicals.” The remaining genes were categorized as “unaffected pathogenicity” (24) and “loss of pathogenicity” (5). Notably, 22 genes related to human diseases (DFVF-HUM) encoded lethal virulence factors, while two encoded hypervirulence factors (Fig. 3D). We also screened the CAZymes, SMs, and pathways related to iron metabolism and stress responses associated with human pathogenicity by comparing their gene profiles with DFVF-HUM (Tables S12 and S13). As shown in Table 6 and Fig. 5A, *C. bantiana* shared five genes primarily linked to chitin synthase and 1,3-β-D-glucan synthase between CAZymes and DFVF-HUM, and four genes associated with PKS, α-tubulin, and actin-like protein 2 between SMs and DFVF-HUM. We also identified four genes related to MAPK, SOD, and catalase-peroxidase that were shared between stress response pathways and DFVF-HUM (Fig. 5B).

**Figure 5.**
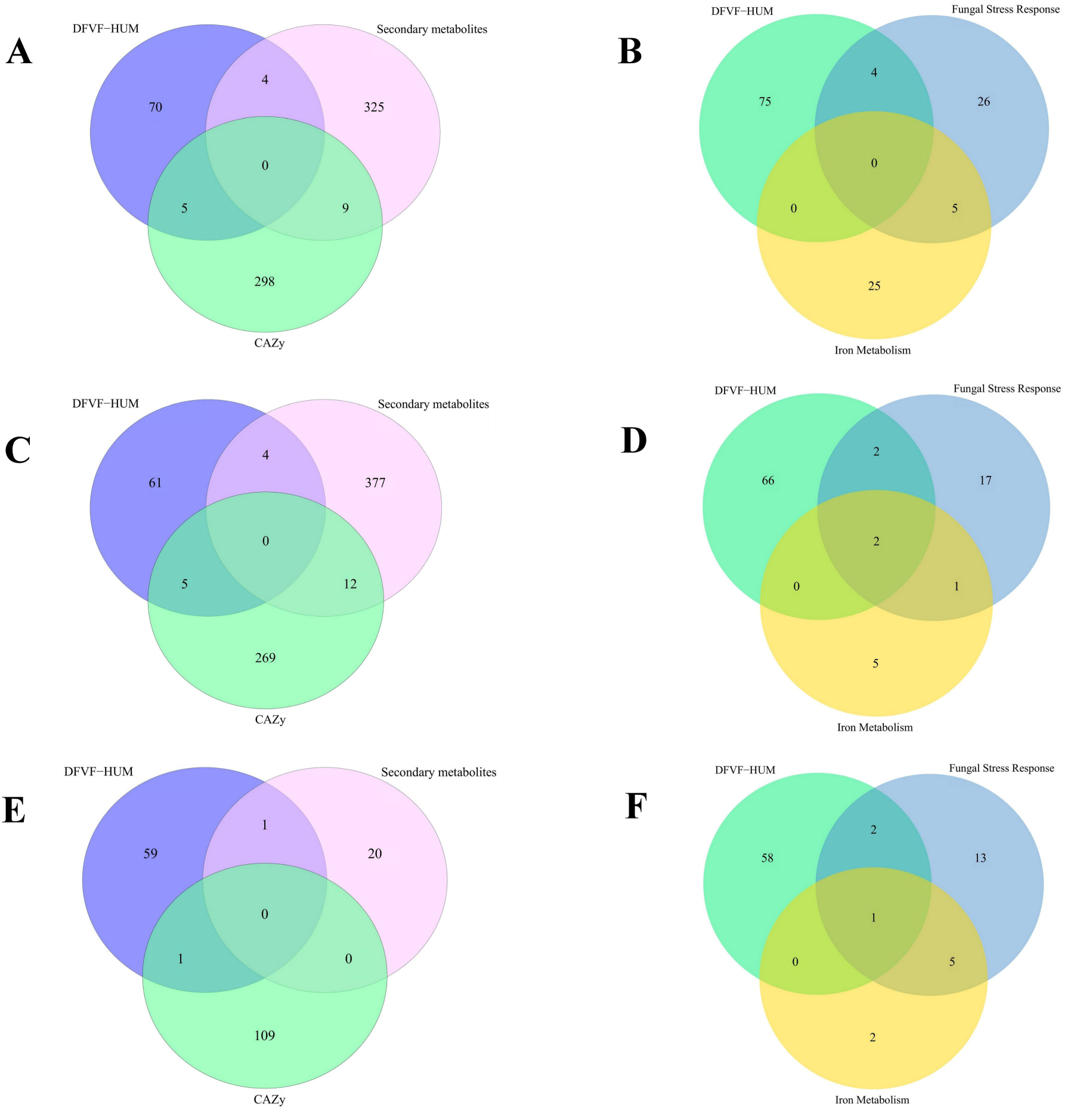
Analysis of genes encoding CAZymes, secondary metabolites, and proteins involved in iron metabolism and stress responses related to human diseases. Venn diagrams illustrate the genes encoding DFVF-HUM, CAZymes, and secondary metabolites in *C. bantiana* (A), *F. monophora* (C), and *C. cladosporioides* (E). Additionally, the genes encoding DFVF-HUM and proteins involved in iron metabolism and stress responses in *C. bantiana* (B), *F. monophora* (D), and *C. cladosporioides* (F) are also shown.

**Table 6.** Genes shared between metabolic pathways and human diseases.

| Species | Shared Gene IDs | Protein encoded |
| --- | --- | --- |
| **Carbohydrate-active enzymes** |  |  |
| *C. bantiana* | NMHCB_012341 | Uncharacterized protein Z519 |
| | NMHCB_002571 | α-1,2-mannosyltransferase |
|  | NMHCB_009723 | Chitin synthase |
| | NMHCB_004231 | 1,3-β-D-glucan synthase |
|  | NMHCB_003333 | Chitin synthase |
| *F. monophora* | NMHFM_011587 | 1,3-β-D-glucan synthase |
| | NMHFM_003258 | α-1,2-mannosyltransferase |
|  | NMHFM_010830 | Chitin synthase |
|  | NMHFM_003591 | Alpha-glucuronidase A |
|  | NMHFM_008789 | Chitin synthase |
| *C. cladosporioides* | NMHCC_008369 | 1,3-β-D-glucan synthase |
| **Secondary metabolites** |  |  |
| *C. bantiana* | NMHCB_006641 | Polyketide synthase |
|  | NMHCB_009559 | Start control protein cdc10 |
|  | NMHCB_007256 | Actin-like protein 2 |
|  | NMHCB_000629 | Tubulin alpha chain |
| *F. monophora* | NMHFM_004562 | Tubulin alpha chain |
|  | NMHFM_006093 | Tryptophan synthase |
|  | NMHFM_001312 | Actin-like protein 2 |
|  | NMHFM_000705 | Hypothetical protein AYO21 03016 |
| *C. cladosporioides* | NMHCC_009445 | GATA transcription factor (White collar 2) |
| **Iron metabolism** |  |  |
| *F. monophora* | NMHFM_005515 | Hog1 MAPK |
|  | NMHFM_011208 | MAPK |
| *C. cladosporioides* | NMHCC_005288 | MAPK |
| **Stress response** |  |  |
| *C. bantiana* | NMHCB_003738 | MAPK |
|  | NMHCB_008065 | MAPK/P38 protein kinase |
|  | NMHCB_007585 | Superoxide dismutase |
|  | NMHCB_005801 | Catalase-peroxidase |
| *F. monophora* | NMHFM_009985 | Superoxide dismutase |
|  | NMHFM_008185 | Catalase-peroxidase |
|  | NMHFM_005515 | Hog1 MAPK |
|  | NMHFM_011208 | MAPK |
| *C. cladosporioides* | NMHCC_000226 | Superoxide dismutase |
|  | NMHCC_005288 | MAPK |
|  | NMHCC_004108 | Hog1 MAPK |

MAPK, mitogen-activated protein kinase; Hog1, high osmolarity glycerol

In *F. monophora*, 66 virulence-related genes were identified, with 64 associated with reduced virulence and two (NMHFM_005262 and NMHFM_011267) linked to hypervirulence. This species exhibited nine “lethal” genes and two associated with “resistance to chemicals.” The remaining genes comprised those classified as “unaffected pathogenicity” (27) and “loss of pathogenicity” (6). As illustrated in Fig. 3E, 22 genes related to human diseases encoded lethal virulence factors, and four encoded hypervirulence factors. *F. monophora* shared five genes primarily linked to chitin synthase and 1,3-β-D-glucan synthase between CAZymes and DFVF-HUM, and four genes associated with tryptophan synthase and actin-like protein 2 between SMs and DFVF-HUM (Table 6 and Fig. 5C). We also identified four genes shared between stress response pathways and DFVF-HUM, including two Hog1 MAPK genes that were shared among iron metabolism, stress response, and DFVF-HUM (Fig. 5D).

For *C. cladosporioides*, 48 virulence-related genes were identified, with 46 linked to reduced virulence and two (NMHCB_003893 and NMHCB_006928) associated with hypervirulence. There were 10 “lethal” genes and one involved in “resistance to chemicals.” The remaining genes were categorized as “unaffected pathogenicity” (22) and “loss of pathogenicity” (10). Analysis of DFVF-HUM indicated that 14 genes encoding lethal virulence factors were associated with human diseases, and four were linked to hypervirulence (Fig. 3F). *C. cladosporioides* shared a gene linked to 1,3-β-D-glucan synthase between CAZymes and DFVF-HUM, as well as a unique GATA transcription factor between SM and DFVF-HUM (Table 6 and Fig. 5E). We also identified three genes, primarily related to Hog 1 MAPK and SOD, that were shared between stress response pathways and DFVF-HUM, with one MAPK gene also linked to iron metabolism (Fig. 5F).

### Antifungal resistance genes and efflux transporters

In *C. bantiana*, annotation with MARDy database revealed a single ERG11/CYP51a gene (NMHCB_009218) encoding lanosterol 14α demethylase, and involved in azole resistance. Additionally, we identified FKS1 (NMHCB_004231) and FKS2 (NMHCB_004231) genes, both encoding 1,3-beta-glucan synthase and associated with echinocandin resistance, as well as with a FUR1 gene (NMHCB_007550) encoding uracil phosphoribosyltransferase (UPRTase) and mediating resistance to 5-flucytosine (5-FC). In *F. monophora*, there were two ERG11/CYP51a genes (NMHFM_004941 and NMHFM_011144), along with a FKS1 (NMHFM_011587) and FKS2 gene (NMHFM_011587), as well as a FUR1 gene (NMHFM_010011). For *C. cladosporioides*, one ERG11/CYP51a gene (NMHCC_010684), a FKS1 (NMHCC_008369) and FKS2 gene (NMHCC_008369), and two FUR1 genes (NMHCC_004903 and NMHCC_002572) were annotated.

Antifungal resistance is often driven by the overexpression of multidrug efflux transporters, including major facilitator superfamily (MFS) and ATP-binding cassette (ABC) transporters. In this study, we identified several MFS transporters in *C. bantiana* and *F. monophora*, with the most prominent being the general substrate transporter (GST), sugar porter (SP), drug:H+ antiporter-1 (DHA1), multidrug resistance protein 1 (MDR1), and anion:cation symporter (ACS). In contrast, *C. cladosporioides* harboured genes for only GST (1) and MDR1 (2) types of MFS transporters. Notably, drug:H+ antiporter-2 (DHA2) and L-amino acid transporter-3 (LAT3) were exclusively found in *C. bantiana*. ABC transporters were present in all three species, with the CDR1 gene being unique to *C. bantiana* (Table 7). Further details on the annotations of multidrug efflux transporter genes are available in Table S14.

**Table 7.** Analysis of multidrug efflux transporters.

| **MFS transporters** | **Number of genes** |  |  |
| --- | --- | --- | --- |
|  | **C. bantiana** | **F. monophora** | **C. cladosporioides** |
| GST | 61 | 19 | 1 |
| SP | 29 | 9 | 0 |
| DHA1 | 23 | 2 | 0 |
| MDR1 | 29 | 3 | 2 |
| ACS | 10 | 3 | 0 |
| MCP | 8 | 1 | 0 |
| FHS | 6 | 4 | 0 |
| DHA2 | 4 | 0 | 0 |
| SHS | 3 | 2 | 0 |
| SIT | 2 | 2 | 0 |
| LAT3 | 1 | 0 | 0 |
| NNP | 1 | 1 | 0 |
| **ABC transporters** | 30 | 42 | 17 |
| **CDR1** | 1 | 0 | 0 |

MFS, major facilitator superfamily; GST, general substrate transporter; SP, sugar porter; DHA1, drug: H<sup>+</sup> antiporter-1; DHA2, drug: H<sup>+</sup> antiporter-2; MDR1, multidrug resistance protein 1; ACS, anion: cation symporter; MCP, monocarboxylate porter; FHS, fucose: H<sup>+</sup> symporter; SHS, sialate: H<sup>+</sup> symporter; SIT, siderophore-iron transporter; LAT3, L-amino acid transporter-3; NNP, nitrate/nitrite transporter; ABC, ATP-binding cassette transporter; CDR1, Candida drug resistance-1

## DISCUSSION

Our study unravels the first reference genome assemblies of three neurotropic dematiaceous fungal species: *C. bantiana*, *F. monophora*, and *C. cladosporioides* using Oxford Nanopore long-read sequencing technology. It offers novel insights into their metabolic capabilities, pathogenic potential, and evolutionary adaptations. By conducting KEGG pathway analysis, GO annotations, and identifying various gene families, we unveil a complex network of functions that facilitate their survival and interactions with their environment, including human hosts.

The KEGG functional annotation revealed that all three species are equipped with extensive metabolic pathways, particularly in carbohydrate, amino acid, and lipid metabolism. The presence of numerous enzymes related to glycolysis, gluconeogenesis, the tricarboxylic acid (TCA) cycle, and other key carbohydrate metabolism pathways suggests that all three species can effectively metabolize a wide range of complex carbohydrates, including sucrose, galactose, fructose, mannose, pyruvate, and starch. A previous study by Kuan *et al*. [12] also highlighted such abundance of enzymes related to carbohydrate, protein, and lipid metabolism in *C. bantiana*. This metabolic versatility not only underscores their adaptability to different ecological niches but also suggests potential roles in bioremediation and nutrient cycling in various environments. Such capabilities likely contribute to their ubiquitous nature and ecological success.

The GO analysis revealed distinct functional gene profiles among the three species, highlighting their ecological and metabolic diversity. *C. bantiana* exhibits significant enrichments in “molecular function,” particularly in ATP and DNA binding activities, emphasizing its diverse metabolic and functional capabilities. *F. monophora* also shows a striking increase in genes related to “cellular component,” indicating a potentially greater structural complexity, particularly in membrane-related functions. This species stands out with its notable enrichment in zinc ion binding, highlighting a possible specialized role in micronutrient homeostasis. In contrast, *C. cladosporioides* displays a more limited number of genes across all categories, suggesting a narrower functional repertoire. The consistent enrichment in “membrane” and “nucleus” across all three species underscores their fundamental importance in cellular functions, while the variations in specific enrichments suggest differing evolutionary pressures and functional adaptations among the species. Overall, our findings reflect the diversity of functional roles and cellular structures across these fungi, which may influence their ecological niches and interactions.

Genomic plasticity allows organisms to adapt and thrive in diverse environments, a vital trait for pathogens seeking to exploit new niches [52]. Transposable elements (TEs) not only facilitate genomic rearrangements but are also believed to play a role in developing adaptive resistance to antifungal agents [53]. In dematiaceous fungi, the amount of TEs can vary significantly. Consistent with the findings of Teixeira *et al*. [54], our research revealed that the dominant retrotransposon in *C. bantiana* and *F. monophora* is long terminal repeat (LTR), particularly of the Gypsy/DIRS1 type. Similar trends have been observed in other dematiaceous fungal pathogens, including *Bipolaris papendorfii* [55], *Sporothrix schenckii* [56], and *Ochroconis mirabilis* [57]. The notable presence of Class I and Class II TEs in *C. bantiana* suggests a dynamic evolutionary history and significant genome expansion, potentially facilitating rapid adaptation to environmental changes. Conversely, the lower TE content and lack of LTRs in *C. cladosporioides* may indicate a more streamlined genomic structure, possibly reflecting a specialized ecological niche that merits further investigation. The specific TEs identified also suggest functional implications, such as facilitating gene rearrangement and generating novel gene combinations that could enhance adaptability and pathogenic potential. Saprophytic fungi produce a diverse range of CAZymes that facilitate the breakdown, biosynthesis, or modification of carbohydrates and glycoconjugates. Our analysis revealed that *C. bantiana* and *F. monophora* had significantly higher numbers of CAZyme families compared to *C. cladosporioides*. Supporting the findings of Kuan *et al*. [12], we observed that majority of CAZyme-encoding genes in all three species were GHs, followed by GTs. While the specific ecological functions of these enzymes remain uncertain, the absence of polysaccharide lyases (PLs) in *C. bantiana* and *F. monophora* indicates a saprophytic lifestyle, as noted by Zhao *et al*. [58]. Although previous studies have highlighted the importance of cell wall-degrading enzymes, especially GHs, in fungal invasion and pathogenicity in plants [59,60], their potential implications in human diseases have not been thoroughly explored. Liu *et al*. [6] identified 54 CAZyme-encoding genes, primarily GHs and GTs, linked to human diseases in *Paraconiothyrium brasiliense*, isolated from a case of orbital cellulitis. Our analysis revealed that CAZymes, especially 1,3-β-D-glucan synthase, were associated with human diseases across all three species. Moreover, chitin synthase and α-mannosyltransferase were identified as key virulence factors in *C. bantiana* and *F. monophora*.

Fungal SMs are natural products essential for colonizing specific ecological niches. [61]. While some of these metabolites can be toxic to humans, others serve as a source of valuable therapeutic agents [62]. Furthermore, SMs are crucial for the pathogenicity, virulence, and progression of fungal diseases. [63]. Despite their diversity, all SMs are synthesized by a limited number of common biosynthetic pathways, classified based on the enzymes involved: polyketides (PKS), non-ribosomal peptides (NRPS), terpenes, and indole alkaloids [61]. In this study, we identified numerous SM biosynthetic gene clusters, including PKS I/III, NRPS, RiPP, and terpenes, as well as compounds from hybrid pathways, particularly in *C. bantiana* and *F. monophora*. This underscores their ability to produce bioactive compounds that could play a crucial role in ecological competition, communication, and survival across various niches. In contrast, *C. cladosporioides* showed fewer gene clusters, indicating a potential evolutionary trade-off or specialization. Notably, we identified a novel SM gene cluster, NAPAA, in *C. cladosporioides*, recognized for its antioxidant and antimicrobial properties.

Dematiaceous fungi are remarkable for their ability to synthesize melanin in their cell walls. Melanin serves as a virulence factor, providing protection against environmental stresses, including UV radiation, oxidative agents, and extreme temperatures [64]. It also binds to hydrolytic enzymes, inhibiting their action on the plasma membrane, and to antifungal drugs, reducing their effectiveness [64,65]. Furthermore, melanin impairs the phagocytic activity of neutrophils and macrophages, facilitating central nervous system invasion. Disruption of melanin production has been shown to significantly reduce virulence in animal models and restrict hyphal growth [66]. These functions may help elucidate the pathogenic potential of dematiaceous fungi, even in immunocompetent hosts. Dematiaceous fungi can produce melanin through distinct pathways: eumelanin via the 1,8-dihydroxynaphthalene (DHN) and 3,4-dihydroxyphenylalanine (DOPA) pathways, and pyomelanins through l-tyrosine degradation pathway [54]. Notable human pathogens that synthesize DHN-melanin include *Aspergillus nidulans*, *A. niger*, *Alternaria alternata*, *C. bantiana*, *Cladosporium* spp., *E. dermatitidis*, *E. jeanselmei*, *Fonsecaea* spp., *Phialophora* spp., and *Neoscytalidium dimidiatum* [67]. Central to this process are polyketide synthases (PKS), which generate DHN-melanin precursors [64]. Our analysis identified several type I PKS gene clusters linked to melanin production in all three species. Notably, we found a novel type III PKS gene cluster (NMHCC_003783) in *C. cladosporioides*. While PKS III genes are well-characterized in plants and bacteria, their roles in fungal biology remains unclear [68]. These genes often reside near those encoding CYPs, MFS transporters, and transcription factors, contributing to the synthesis of SMs with diverse biological activities, including antimicrobial properties and involvement in disease progression [69]. However, their specific contributions to human diseases warrant further investigation. Consistent with previous reports [64,66], our analysis indicated that SMs, particularly PKS, play a role in the pathogenicity of *C. bantiana* and *F. monophora*. While actin and actin-related proteins have been linked to virulence in phytopathogenic fungi [70], their involvement in human diseases remains largely unexplored. Here, we identified for the first time that genes encoding actin-like protein 2 are associated with human diseases. We also discovered a GATA transcription factor (white collar 2) linked to virulence in *C. cladosporioides*.

Iron is crucial for the survival of nearly all organisms, including fungi, and its acquisition is critical for pathogenicity. Fungi typically employ two main strategies for iron uptake: nonribosomally synthesized secreted iron chelators known as siderophores and reductive iron assimilation (RIA) [71]. Siderophore iron transporters (SITs) are SMs that serve as important virulence factors, being activated in response to interactions with immune cells during infection [72]. In this study, we identified several genes related to iron metabolism in *C. bantiana* and *F. monophora*, including L-ornithine-N_5_-monooxygenase, which is involved in the production of extracellular fusarinine C (FSC) and triacetylfusarinine C, as well as intracellular ferricrocin. We also found SITs and ABC multidrug transporters within the FSC biosynthetic gene cluster, indicating that these fungi have developed robust mechanisms for iron acquisition, crucial for their survival in iron-deficient environments during infection. Additionally, the RIA pathway, which utilizes membrane-bound metalloreductases to convert ferric iron to ferrous iron for cellular uptake, was identified for the first time in *C. bantiana*. The presence of cupin domain-containing proteins further illustrates the versatility of *C. bantiana* in utilizing iron. In contrast, *C*. *cladosporioides* has fewer genes associated with iron metabolism, lacking both SITs and L-ornithine-N_5_-monooxygenase, suggesting it may rely on alternative metabolic pathways or adaptations for iron homeostasis. Notably, the identification of a unique high-affinity iron permease in *C. cladosporioides* indicates species-specific adaptations that merit further research.

Pathogenic fungi often possess a variety of stress-responsive proteins that enhance their pathogenicity and facilitate adaptation to diverse host environments. In this study, we found that all three species harbor essential genes, such as HSP60 and HSF1, which are crucial for survival within the human host. This thermotolerance is a key virulence factor that contributes to their pathogenic potential [73]. Upon experiencing acute heat shock, HSF1 gets phosphorylated and triggers the expression of HSP genes through canonical heat shock elements in their promoters. This induction of HSP genes helps refold or degrade damaged proteins, aiding cellular adaptation to thermal stress [74,75]. Consequently, the HSF1-HSE regulatory network plays a vital role in helping these organisms adapt to thermal stress.

Human phagocytic cells generate ROS, such as superoxide anion (O2−), hydrogen peroxide (H_2_O_2_), and hydroxyl radical (OH), to combat fungal pathogens by damaging their DNA, proteins, and membranes [76]. In turn, fungi have evolved detoxification mechanisms to neutralize these harmful substances. Notably, ROS stress responses can enhance fungal virulence [77]. In our study, we identified several genes associated with ROS detoxification, including SOD, Hog1 MAPK, glutathione peroxidase, monothiol glutaredoxin, and catalases across all three species. This suggests a robust defence system against oxidative damage that supports their survival and pathogenicity. Notably, all three species share stress response genes with human disease-related pathways (DFVF-HUM). Additionally, genes encoding methionine sulfoxide reductase and a ZCTF were exclusively identified in *C. cladosporioides*, which may contribute to its distinct virulence profile. The ZCTF has been shown to mediate core stress responses in *Saccharomyces cerevisiae* and *C. glabrata* [78]. This indicates that while these fungi share common stress adaptation strategies, they also possess specialized mechanisms tailored to their ecological niches and interactions with the host.

The analysis of pathogenicity-associated genes across *C. bantiana*, *F. monophora*, and *C. cladosporioides* reveals intriguing insights into their virulence profiles and potential implications for human health. *C. bantiana* exhibited the highest number of virulence-related genes among the three species, including two associated with hypervirulence. Notably, both *F. monophora* and *C. cladosporioides* also exhibited hypervirulence traits, although they exhibited a predominance of genes linked to reduced virulence. The identification of lethal genes, particularly those related to human diseases, underscores the potential health risks posed by these pathogens. Furthermore, the shared genes between MAPK-mediated stress response pathways and human diseases suggest common virulence mechanisms, indicating a conserved evolutionary strategy for overcoming host defences. This complex interplay of genes represents a promising avenue for further research, as they may significantly influence pathogenicity and drug resistance.

Antifungal therapy in PCP primarily relies on triazoles, either alone or in combination with echinocandins and 5-FC [79]. Dematiaceous fungi show varying susceptibility to antifungal agents, with some species displaying high minimum inhibitory concentrations (MICs) for azoles and 5-FC [80,81]. There is substantial evidence pointing to genetic factors contributing to azole resistance in fungi. This resistance can result from mutations in the enzyme lanosterol 14α demethylase, encoded by the ERG11/CYP51a gene, or from the overexpression of membrane transporters [82]. In particular, ABC and MFS transporters are associated with clinical azole resistance in various *Candida* species [83]. For instance, the upregulation of the major facilitator MDR1 allows *C. albicans* to not only resist azoles but also evade host defences through the efflux of antimicrobial peptides, such as histatin 5 [84]. Echinocandin resistance, on the other hand, is linked to specific mutations in the hotspot regions of the FKS gene, which encodes 1,3-β-D-glucan synthase, the target for echinocandins. Furthermore, resistance and tolerance to echinocandins can be influenced by complex cellular mechanisms that regulate responses to cell wall stress, such as the upregulation of chitin production mediated by calcineurin, protein kinase C, the Hog1 MAPK pathway, and the molecular chaperone Hsp90 [85]. Resistance to 5-FC in clinically relevant fungal species typically results from mutations in genes like FCY2, FCY1, FUR1, and UXS1, which are involved in cytosine transport and metabolism [86]. Our study identified several genes linked to antifungal resistance across all three species, including ERG11/CYP51a for azoles, FKS1 and FKS2 for echinocandins, and FUR1 for 5-FC. The unique efflux transporter profiles indicate varied resistance mechanisms, with *C. bantiana* exhibiting a broader array of transporters that may confer a survival advantage in antifungal-rich milieu. This underscores the potential for treatment failure, complicating the management of such cases. Further research into the genetic and cellular factors driving these mechanisms is crucial for improving our understanding of antifungal resistance in these species.

## CONCLUSIONS

The genomic analyses of *C. bantiana*, *F. monophora*, and *C. cladosporioides* provide valuable insights into their genetic architecture and functional capabilities. The use of long-read sequencing technology facilitated the assembly of highly complete genomes, revealing diverse metabolic pathways, virulence factors, and antifungal resistance mechanisms. Their metabolic versatility, evidenced by a rich repertoire of carbohydrate-active enzymes and secondary metabolite biosynthetic gene clusters, further underscores their ecological adaptability. Notably, the presence of numerous pathogenicity-related genes in all three species raises critical concerns for human health. Overall, these findings enhance our understanding of their roles in both environmental and clinical contexts. The virulence factors identified may serve as potential targets for drug development and vaccine candidates, paving the way for novel therapeutic strategies to improve the management of infections caused by these pathogens.

## ACKNOWLEDGEMENTS

This study was funded by the Indian Council of Medical Research (ICMR), New Delhi, under Grant No. Myco/Adhoc/2/2022-ECD-II. The contents of this publication are the sole responsibility of the authors and do not necessarily reflect the views of the ICMR. The funding agency had no role in study design, data collection and analysis, or preparation of the manuscript. We acknowledge the support of Anuraj. O.P. from Molsys Private Limited (Bengaluru, Karnataka, India) for whole-genome sequencing and genomic data analysis. A.S. contributed to the study design and conceptualization. A.S. and J.M.M. contributed to laboratory data generation. A.S. contributed to data analysis, report writing, and editing. N.S., U.P., H.K., S.M.R., and A.J.K. contributed to report editing and feedback. A.S. contributed to the illustration design. All authors have read and approved the final version of the manuscript prior to submission for publication.

## DATA AVAILABILITY

The whole-genome sequencing data and annotated genome assemblies from this study were submitted to GenBank under BioProject accession numbers [PRJNA1170874](https://www.ncbi.nlm.nih.gov/bioproject/1170874), [PRJNA1170878](https://www.ncbi.nlm.nih.gov/bioproject/1170878), and [PRJNA1170879](https://www.ncbi.nlm.nih.gov/bioproject/1170879). The partial gene sequences are available in GenBank under accession numbers PP462151, PP462152, and PP462153.

