## Supplemental tables S1- S14, Supplemental figures S1-S4 for "Long-read whole-genome sequencing offers novel insights into the biology, stress adaptation, and virulence of neurotropic dematiaceous fungi associated with primary cerebral phaeohyphomycosis": Supplemental Material Legends.docx

**Tables SI, S2, S4, S6, S10; Fig. S1 to S4.** Analysis of KEGG and GO functional annotations, CAZymes, peptidases, iron metabolism, and secondary metabolites in *C. bantiana*, *F. monophora*, and *C. cladosporioides*.

**Table S3.** KEGG and GO functional annotations for *C. bantiana*, *F. monophora*, and *C. cladosporioides*

**Table S5.** CAZyme class annotation distributions in *C. bantiana*, *F. monophora*, and *C. cladosporioides*

**Table S7.** Annotation of genes encoding Cytochrome P450 using FCPD

**Table S8.** Annotation of genes encoding transmembrane proteins using TMHMM

**Table S9.** Annotation of secondary metabolite biosynthetic gene clusters

**Table S11.** Annotation of pathogenicity-associated genes using PHI-base

**Table S12.** List of genes shared between iron metabolism pathways and DFVF-HUM

**Table S13.** List of genes shared between stress response pathways and DFVF-HUM

**Table S14.** Annotation of genes encoding multidrug efflux transporters
