## Supplemental tables S1- S14, Supplemental figures S1-S4 for "Long-read whole-genome sequencing offers novel insights into the biology, stress adaptation, and virulence of neurotropic dematiaceous fungi associated with primary cerebral phaeohyphomycosis": Supplemental material.docx

**Table S1. Major KEGG metabolic pathways of *C. bantiana*, *F. monophora,* and *C. cladosporioides***

| **KEGG Pathways** | **No. of genes** | | |
| --- | --- | --- | --- |
|  | **C. bantiana** | **F. monophora** | **C. cladosporioides** |
| Metabolic pathways | 1198 | 1264 | 1179 |
| Biosynthesis of secondary metabolites | 508 | 537 | 476 |
| Microbial metabolism in diverse environments | 368 | 380 | 294 |
| Carbon metabolism | 149 | 155 | 136 |
| Biosynthesis of cofactors | 149 | 174 | 145 |
| Biosynthesis of amino acids | 130 | 124 | 129 |
| Starch and sucrose metabolism | 0 | 0 | 91 |
| Tyrosine metabolism | 84 | 78 | 0 |
| Pyruvate metabolism | 84 | 90 | 56 |
| Chemical carcinogenesis- reactive oxygen species | 82 | 81 | 83 |
| Glycolysis / Gluconeogenesis | 77 | 86 | 55 |
| Valine, leucine and isoleucine degradation | 73 | 97 | 51 |
| Oxidative phosphorylation | 65 | 62 | 73 |
| Tryptophan metabolism | 65 | 73 | 60 |
| Fatty acid metabolism | 64 | 75 | 0 |
| Butonate metabolism | 0 | 61 | 0 |
| Arginine and proline metabolism | 60 | 73 | 59 |
| Amino sugar and nucleotide sugar metabolism | 0 | 0 | 59 |
| Fatty acid degradation | 59 | 80 | 0 |
| Degradation of aromatic compounds | 59 | 79 | 0 |
| Peroxisome | 56 | 67 | 0 |
| Purine metabolism | 56 | 56 | 52 |
| Glycine, serine and threonine metabolism | 54 | 57 | 51 |
| Lysine degradation | 0 | 54 | 0 |
| Oxocarboxylic acid metabolism | 52 | 0 | 0 |
| beta-Alanine metabolism | 52 | 65 | 0 |
| Drug metabolism- cytochrome P450 | 51 | 0 | 0 |
| Cysteine and methionine metabolism | 50 | 54 | 50 |
| Glycerolipid metabolism | 0 | 53 | 0 |
| Thermogenesis | 0 | 0 | 54 |
| Amyotrophic lateral sclerosis | 49 | 0 | 54 |
| Pathways of neurodegeneration- multiple diseases | 0 | 0 | 54 |
| Diabetic cardiomyopathy | 0 | 0 | 53 |
| Parkinson’s disease | 0 | 0 | 50 |
| Pentose and glucuronate interconversions | 0 | 0 | 49 |
| Alzheimer disease | 0 | 0 | 49 |

**Table S2. Major KEGG pathways and the number of genes associated with human infections in the three species**

| **KEGG pathway** | **Human infections** | **Number of genes** | | |
| --- | --- | --- | --- | --- |
|  |  | **C. bantiana** | **F. monophora** | **C. cladosporioides** |
| KO:05165 | HPV infection | 14 | 12 | 12 |
| KO:05110 | *Vibrio cholerae* infection | 12 | 11 | 11 |
| KO:05120 | *Helicobacter pylori* infection | 12 | 11 | 11 |
| KO:05152 | Tuberculosis | 9 | 7 | 9 |
| KO:05131 | Shigellosis | 7 | 8 | 8 |
| KO:05132 | *Salmonella* infection | 3 | 3 | 3 |
| KO:05143 | African trypanosomiasis | 3 | 4 | 3 |
| KO:05166 | HTLV 1 infection | 2 | 1 | 1 |
| KO:05164 | Influenza A | 2 | 2 | 2 |
| KO:05167 | KSHV infection | 2 | 2 | 2 |
| KO:05130 | Escherichia coli infection | 2 | 2 | 2 |
| KO:05170 | HIV 1 infection | 1 | 1 | 1 |
| KO:05161 | Hepatitis B infection | 1 | 1 | 1 |
| KO:05160 | Hepatitis C infection | 1 | 1 | 1 |
| KO:05162 | Measles | 1 | 1 | 1 |
| KO:05168 | HSV 1 infection | 1 | 1 | 1 |
| KO:05163 | CMV infection | 1 | 1 | 1 |
| KO:05169 | EBV infection | 1 | 1 | 1 |
| KO:05135 | Yersinia infection | 1 | 1 | 1 |
| KO:05134 | Legionellosis | 1 | 1 | 1 |
| KO:05146 | Amoebiasis | 1 | 1 | 0 |
| KO:05145 | Toxoplasmosis | 1 | 1 | 1 |

HPV, human papillomavirus; HTLV 1, human T-cell lymphotropic virus type 1; KSHV, Kaposi sarcoma-associated herpesvirus; HIV 1, human immunodeficiency virus type 1; HSV 1, herpes simplex virus type 1; CMV, cytomegalovirus; EBV, Epstein-Barr virus

**Table S4. Analysis of carbohydrate-active enzymes**

| **Enzyme category** | **C. bantiana** | | **F. monophora** | | **C. cladosporioides** | |
| --- | --- | --- | --- | --- | --- | --- |
|  | **Genes** | **Families** | **Genes** | **Families** | **Genes** | **Families** |
| Glycoside hydrolases | 141 | 56 | 123 | 54 | 53 | 32 |
| Glycosyltransferases | 91 | 30 | 89 | 30 | 26 | 14 |
| Carbohydrate esterases | 6 | 3 | 8 | 4 | 5 | 3 |
| CBMs | 11 | 6 | 12 | 8 | 7 | 4 |
| Polysaccharide lyases | 0 | 0 | 0 | 0 | 7 | 3 |
| Auxiliary activities | 49 | 13 | 43 | 14 | 16 | 8 |

CBM, carbohydrate-binding motif

**Table S6. Analysis of extracellular peptidases**

| **Enzyme category** | **Number of genes** | | |
| --- | --- | --- | --- |
|  | **C. bantiana** | **F. monophora** | **C. cladosporioides** |
| Serine peptidase | 60 | 62 | 8 |
| Aspartic peptidase | 15 | 15 | 1 |
| Metallopeptidase | 51 | 52 | 9 |
| Cysteine peptidase | 47 | 44 | 3 |
| Threonine peptidase | 19 | 18 | 11 |
| Glutamic peptidase | 1 | 1 | 0 |

**Table S10. Analysis of genes involved in iron uptake and homeostasis**

| **Predicted proteins** | **Number of genes** | | |
| --- | --- | --- | --- |
|  | **C. bantiana** | **F. monophora** | **C. cladosporioides** |
| Siderophore iron transporter | 8 | 2 | 0 |
| Cupin domain-containing protein | 7 | 0 | 0 |
| Metalloreductase | 4 | 0 | 0 |
| pH response regulator | 3 | 0 | 2 |
| Vacuolar transporter | 3 | 1 | 1 |
| Mitogen-activated protein kinase | 2 | 3 | 3 |
| Arginase | 1 | 1 | 0 |
| L-ornithine-N^5^-monooxygenase | 1 | 1 | 0 |
| Sterol regulatory element binding protein | 1 | 0 | 0 |
| Zinc cluster transcription factor | 0 | 0 | 1 |
| High affinity iron permease | 0 | 0 | 1 |

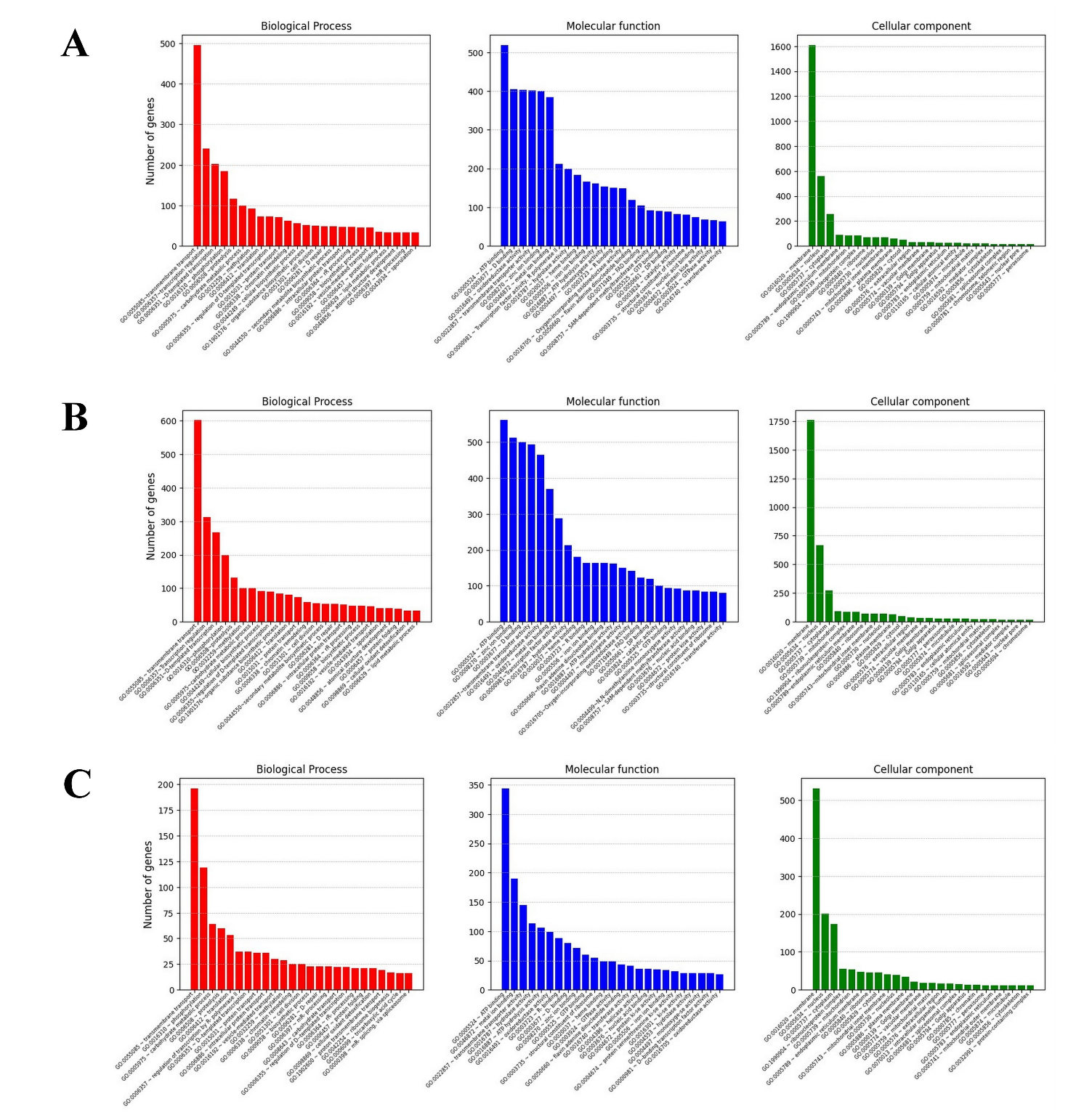

**Figure S1.** Gene Ontology (GO) functional annotations of *C. bantiana* **(A)**, *F. monophora* **(B)**, and *C. cladosporioides* **(C)** genomes. In each category, the top 25 GO terms ranked based on gene counts are shown. The GO enrichment analysis was performed using PANNZER2 web server.

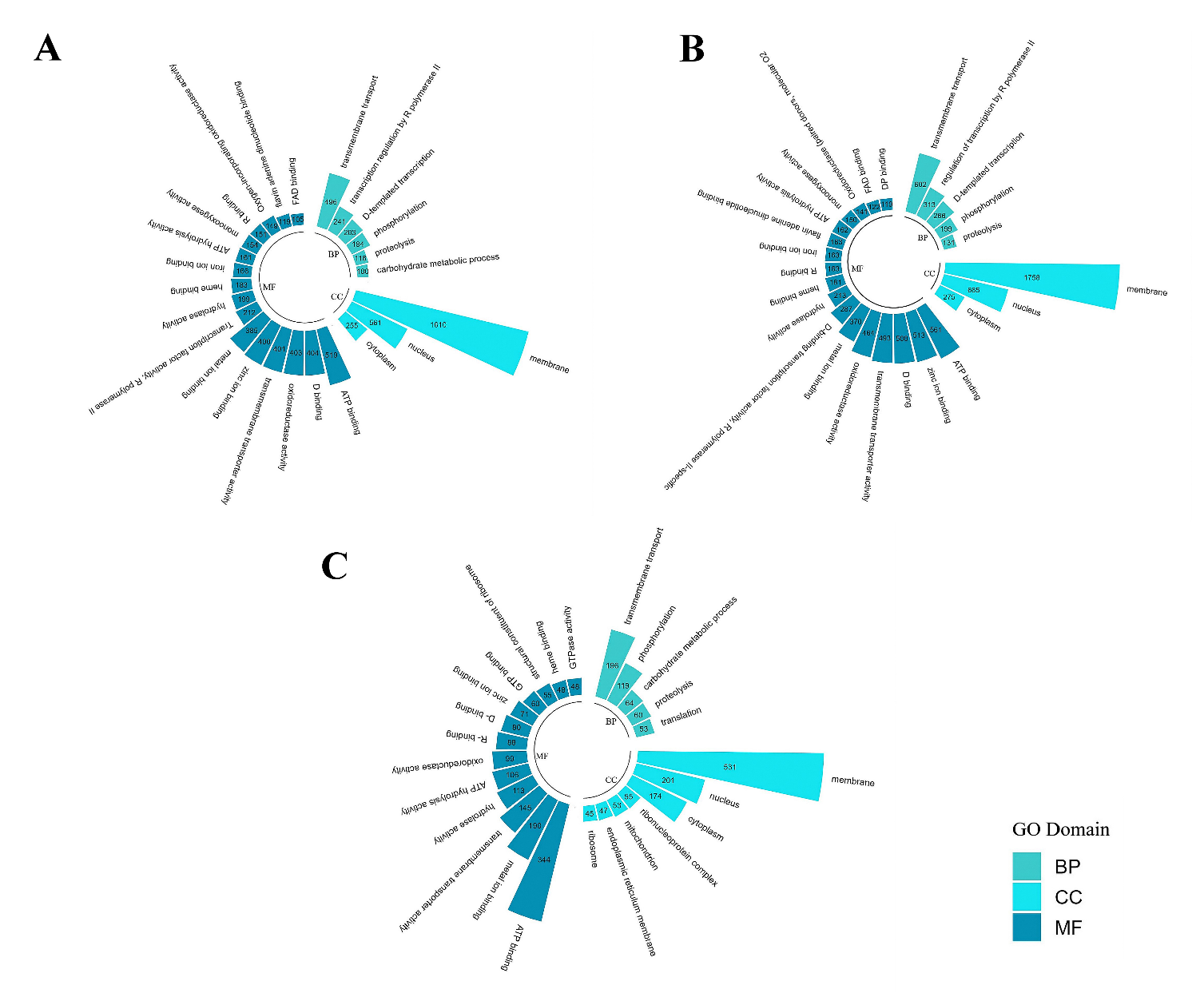

**Figure S2.** Circular Gene Ontology (GO) plot showing functional annotations of *C. bantiana* **(A)**, *F. monophora* **(B)**, and *C. cladosporioides* **(C)** genomes, across three categories: biological processes (BP), cellular components (CC), and molecular functions (MF). The top 25 GO terms ranked based on gene counts are shown. The GO enrichment analysis was performed using PANNZER2 web server.

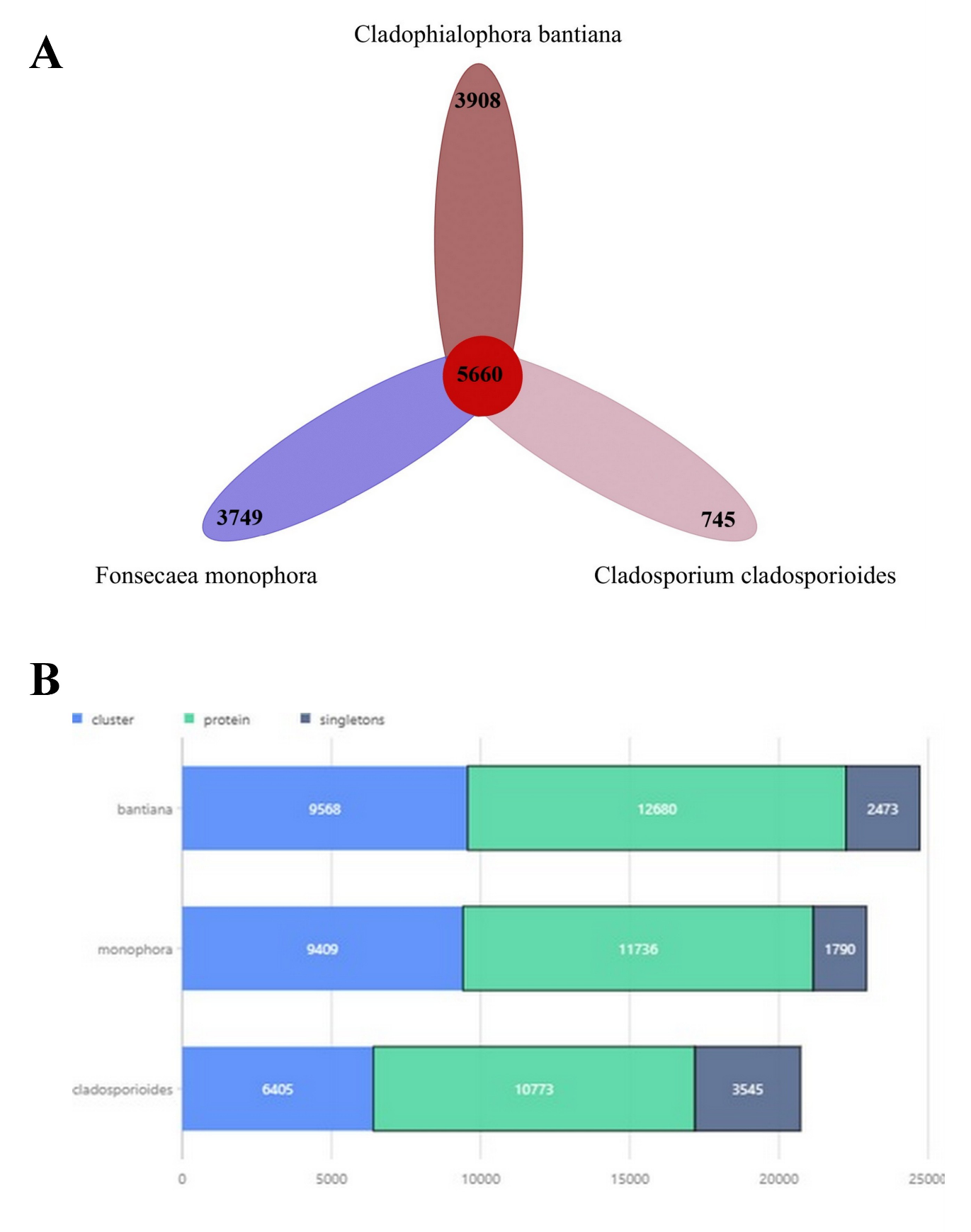

**Figure S3.** Comparative analysis of protein sequences between *C. bantiana*, *F. monophora*, and *C. cladosporioides*, using OrthoVenn3. **A.** Flowerplot showing shared and unique genes among three species. **B.** Composite barplot showing orthologous clusters, unique sequences and singletons in three species.

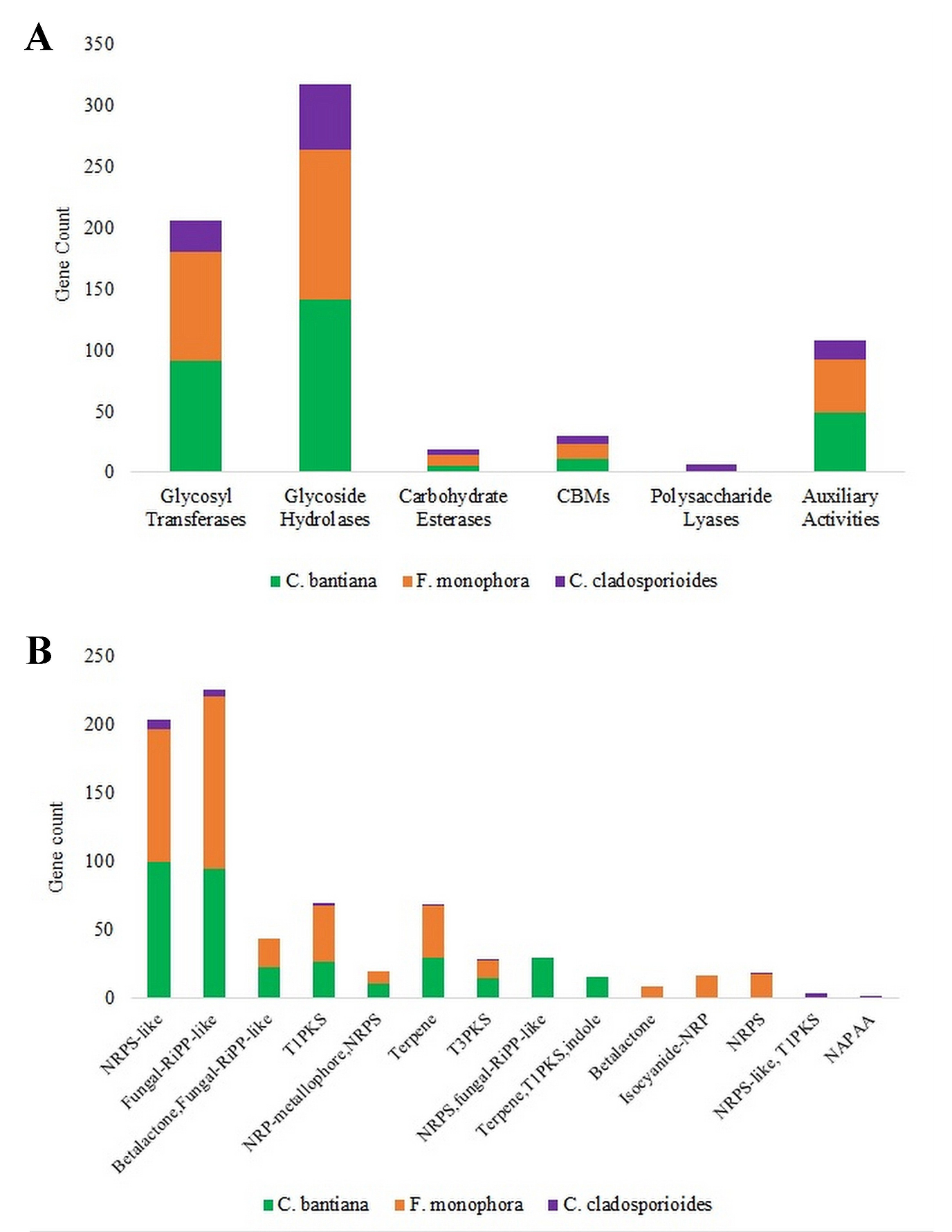

**Figure S4.** Analysis of carbohydrate-active enzymes **(A)** and secondary metabolite biosynthetic gene clusters **(B)** in *C. bantiana*, *F. monophora*, and *C. cladosporioides*. Abbreviations: CBM, carbohydrate-binding motif; NRP, Nonribosomal peptide; NRPS, Nonribosomal peptide synthetase; RiPP, Ribosomally synthesized and post-translationally modified peptides; T1PKS, Type I polyketide synthase; T3PKS, Type III polyketide synthase; NAPAA, non-alpha poly-amino acid
