## Supplemental tables S1- S14, Supplemental figures S1-S4 for "Long-read whole-genome sequencing offers novel insights into the biology, stress adaptation, and virulence of neurotropic dematiaceous fungi associated with primary cerebral phaeohyphomycosis": Supplemental material.pdf

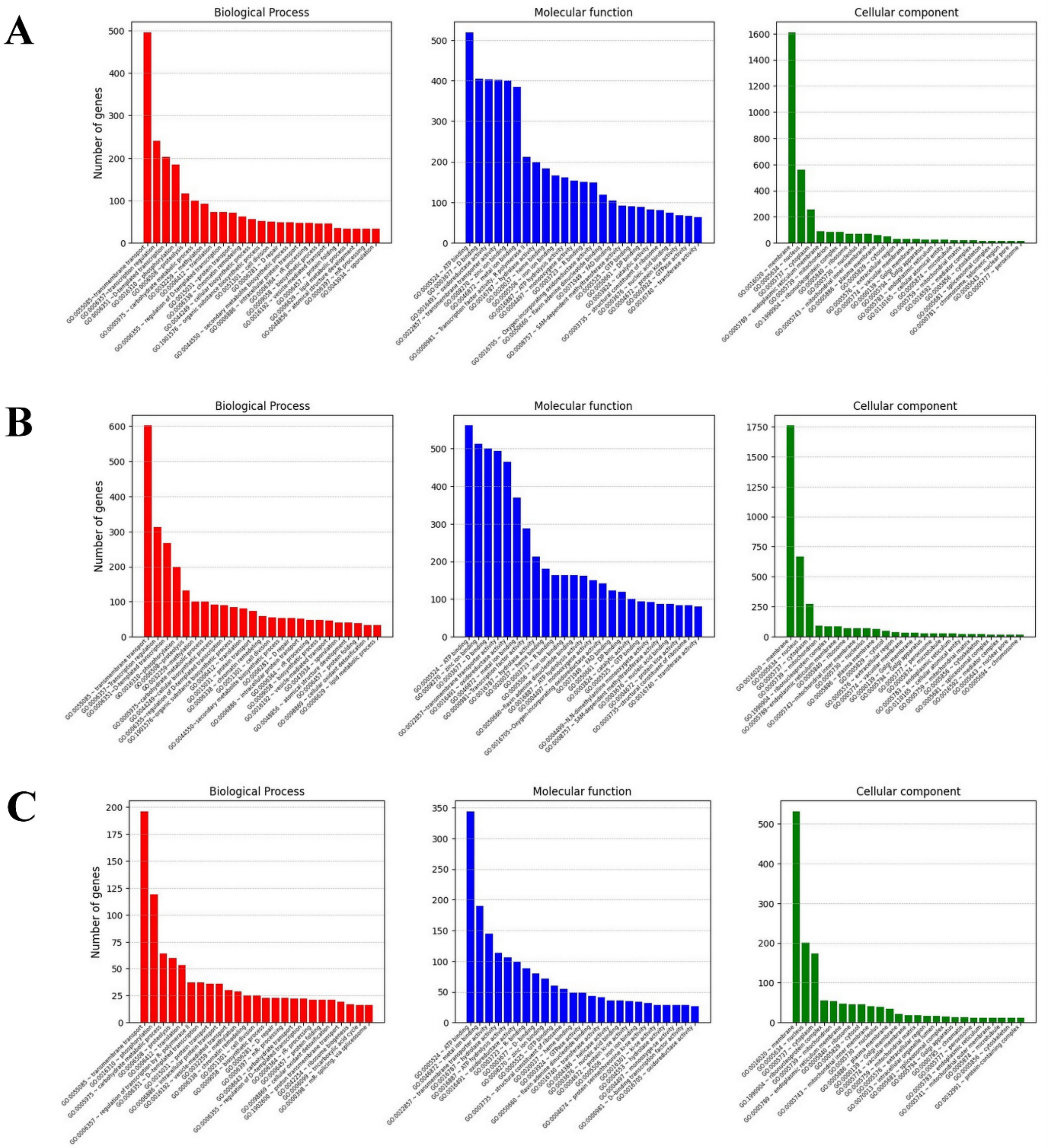


**Figure S1.** Gene Ontology (GO) functional annotations of *C. bantiana* **(A)**, *F. monophora* **(B)**, and *C. cladosporioides* **(C)** genomes. In each category, the top 25 GO terms ranked based on gene counts are shown. The GO enrichment analysis was performed using PANNZER2 web server.

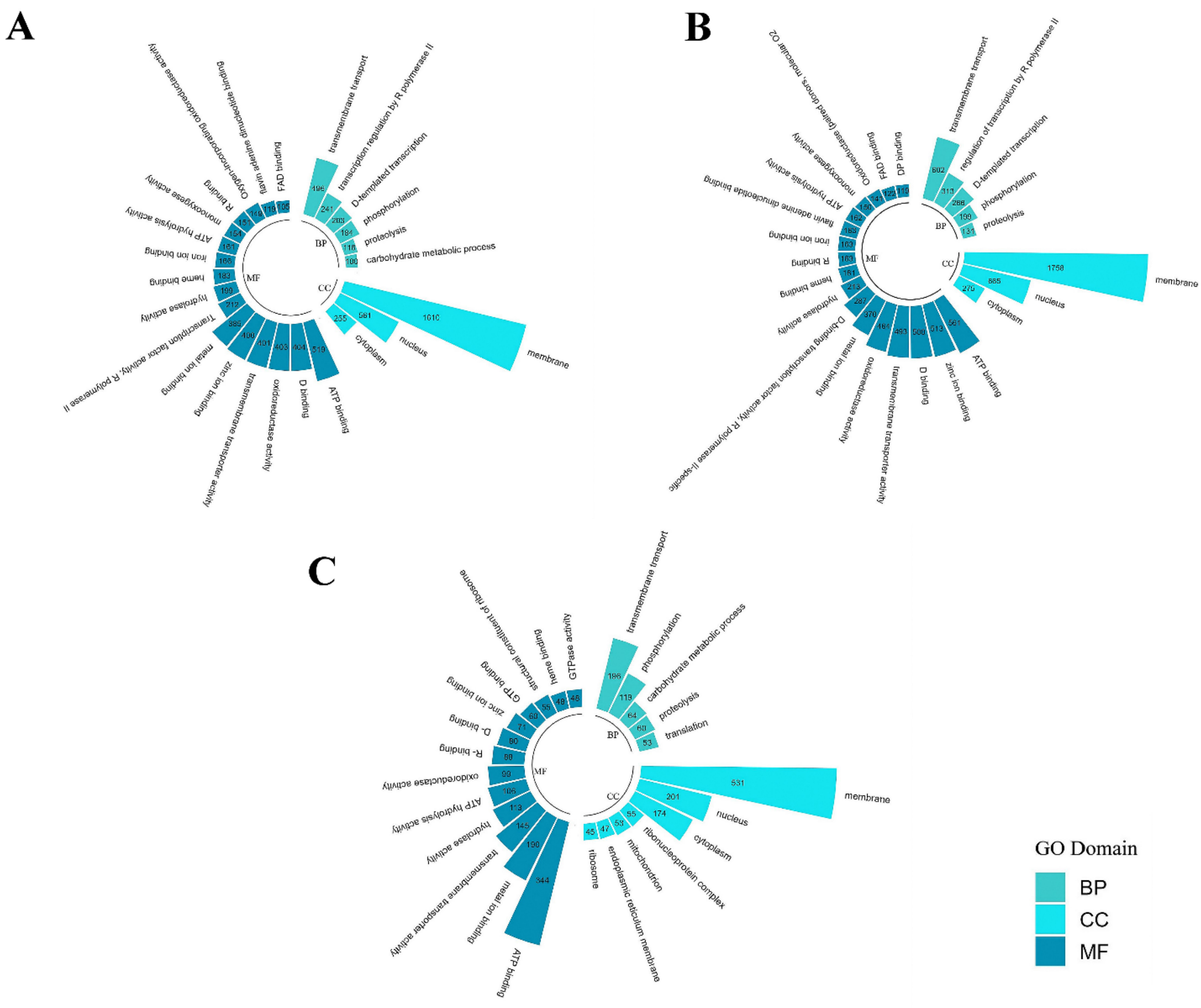


**Figure S2.** Circular Gene Ontology (GO) plot showing functional annotations of *C. bantiana* **(A)**, *F. monophora* **(B)**, and *C. cladosporioides* **(C)** genomes, across three categories: biological processes (BP), cellular components (CC), and molecular functions (MF). The top 25 GO terms ranked based on gene counts are shown. The GO enrichment analysis was performed using PANNZER2 web server.

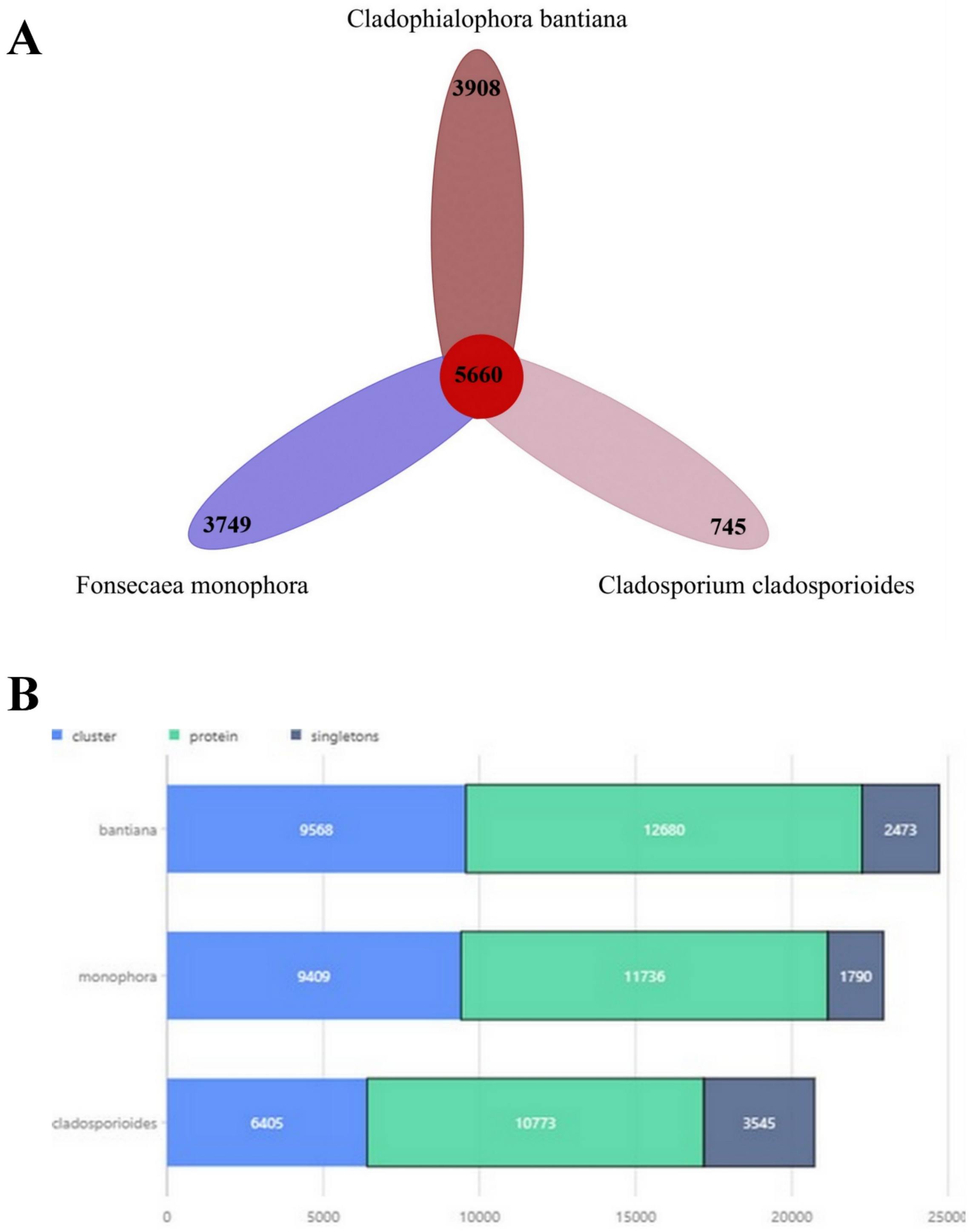


**Figure S3.** Comparative analysis of protein sequences between *C. bantiana*, *F. monophora*, and *C. cladosporioides*, using OrthoVenn3. **A.** Flowerplot showing shared and unique genes among three species. **B.** Composite barplot showing orthologous clusters, unique sequences and singletons in three species.

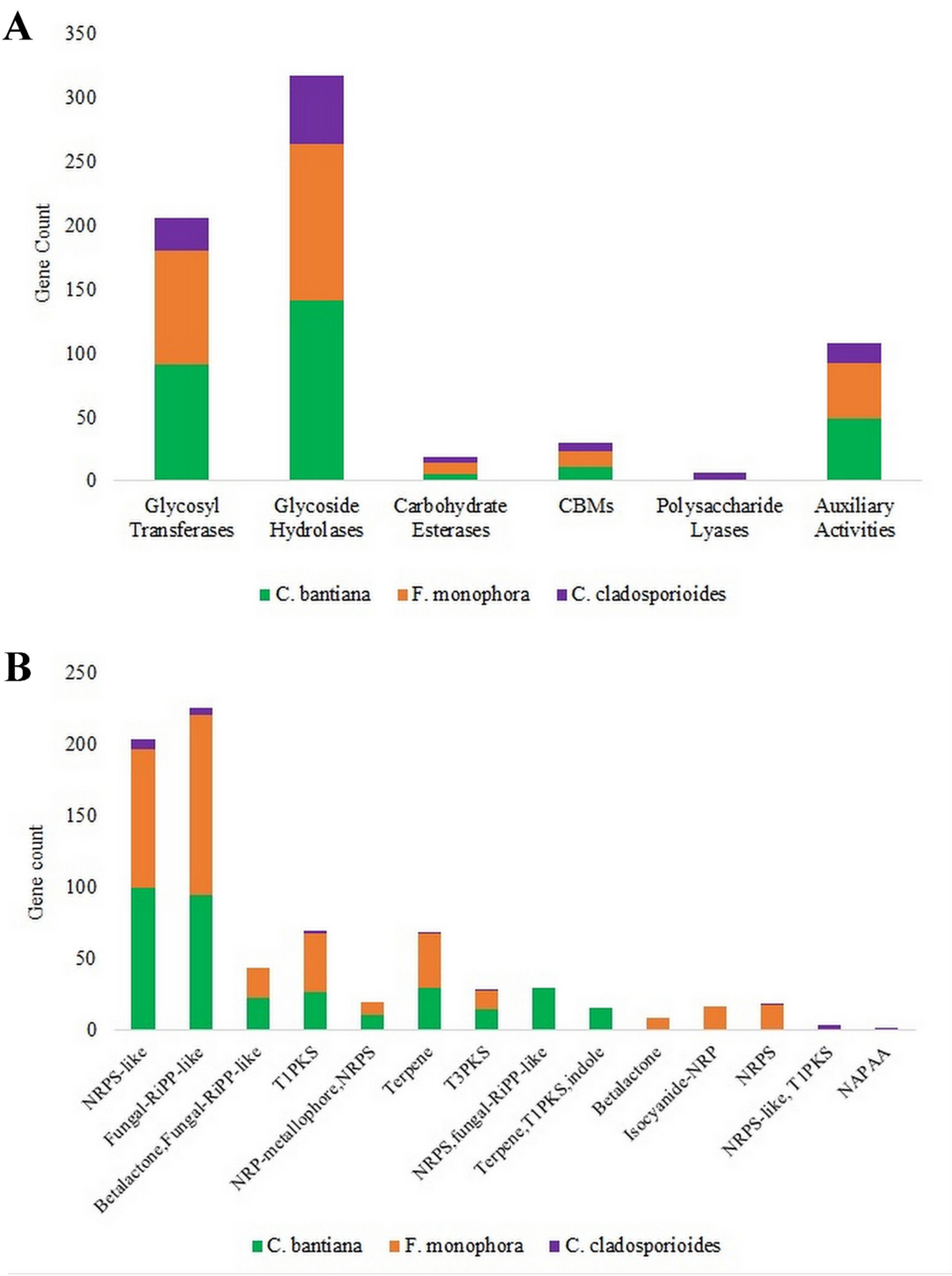


**Figure S4.** Analysis of carbohydrate-active enzymes **(A)** and secondary metabolite biosynthetic gene clusters **(B)** in *C. bantiana*, *F. monophora*, and *C. cladosporioides*. Abbreviations: CBM, carbohydrate-binding motif; NRP, Nonribosomal peptide; NRPS, Nonribosomal peptide synthetase; RiPP, Ribosomally synthesized and post-translationally modified peptides; T1PKS, Type I polyketide synthase; T3PKS, Type III polyketide synthase; NAPAA, non-alpha poly-amino acid
